# Dissecting the genetic determinants of bacterial DNA degradation by bacteriophage T5

**DOI:** 10.64898/2026.04.02.716235

**Authors:** Godfred Morkeyile Annor, Luis Ramirez-Chamorro, Léo Zangelmi, Pascale Boulanger, Ombeline Rossier

**Author notes:** Both authors contributed equally to this work (Godfred Morkeyile Annor, Luis Ramirez-Chamorro).

## Abstract

Upon infection of *Escherichia coli*, the virulent bacteriophage T5 employs a unique two-step mechanism to transfer its 122-kb genome: only 8% of the DNA are initially transferred, allowing expression of pre-early genes that alter host functions before the remaining DNA is delivered. Early infection triggers rapid host DNA degradation and nucleotide catabolism, processes partly controlled by the pre-early genes encoding the predicted metallo-phosphatase A1 and the dNMP phosphatase Dmp. However, the functions of most of the 17 proteins encoded by the first-step transfer DNA (FST-DNA) remain unknown.

Using reverse genetics, we engineered several T5 mutants carrying deletions in the FST-DNA. One of them carries only four pre-early genes (*A1*, *A2*, *hdi* and *009*), demonstrating that thirteen of the 17 predicted genes are dispensable under laboratory conditions. Mutant characterization showed that only *A1* and the predicted DNA-binding protein gene *A2* are essential for productive infection, while *dmp* enhances phage virulence. Notably, seven pre-early genes (*dmp*, *A1*, *hdi*, *hegG*, *011*, *013* and *015*) proved toxic when expressed in *E. coli* without other viral factors, causing severe morphological changes or nucleoid disorganization, while one gene compromised membrane integrity. The essential gene *A1*, which is conserved among all viruses in the *Demerecviridae* family, emerged as the primary driver of host genome degradation, both essential and sufficient for chromosomal DNA digest *in vivo*.

These findings advance our understanding of how T5 manipulates its host during the critical early stages of infection, offering new insights into phage-host interactions and the molecular strategies viruses use to subvert bacterial cells.

## Introduction

Bacteriophages, viruses that infect bacteria, have been studied for over a century and have become indispensable tools in biocontrol, biotechnology, and basic bacterial research. Despite extensive characterization, the functions of many bacteriophage genes remain unknown, even in well-established models such as T4 and T7, which were discovered in the mid-20th century. These uncharacterized genes may play critical roles in manipulating host cellular machinery during early infection. Elucidating their function is crucial not only for advancing our knowledge of phage-host interactions but also for unlocking new applications.

Phage T5, a member of the virulent T-series phages first isolated in 1945, infects *Escherichia coli* and is distinguished by its original two-step DNA transfer mechanism and rapid degradation of host DNA (1, 2). Its 121,750-bp linear double-stranded DNA genome contains two large terminal repeats: the Left Terminal Repeat (LTR) and Right Terminal Repeat (RTR), each spanning 10.2 kb (Fig. 1). The genome is encased within an icosahedral capsid connected to a flexible non-contractile tail. T5 initiates infection by binding to the outer membrane receptor FhuA via the receptor-binding protein pb5, triggering conformational changes in the tail that facilitate DNA delivery across the bacterial envelope (3, 4).

**Figure 1.**
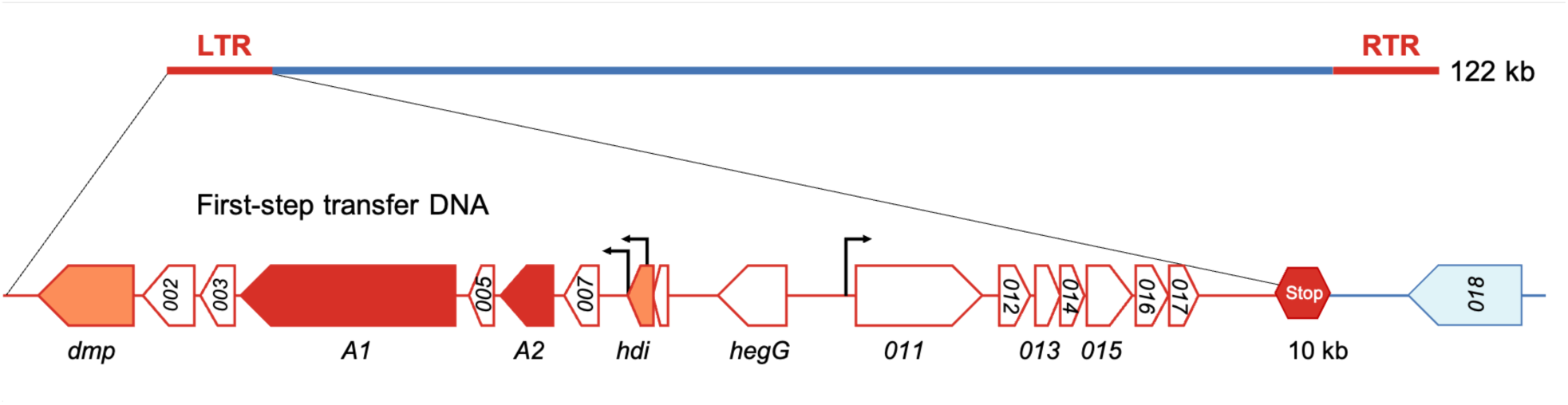
Genetic organization of phage T5 pre-early genes in the Left Terminal Repeat (LTR). The LTR region of the 122-kb T5 genome contains 17 predicted open reading frames (ORFs), with arrows indicating transcription direction (NCBI Reference Sequence: NC_005859.1). An identical gene organization is present in the Right Terminal Repeat (RTR). The First-Step DNA transfer halts after 8% of the genome are delivered to the host cell, as indicated by the stop sign. Among the pre-early genes, *A1* and *A2* (red) are essential for T5 infection: both are required for the Second-Step Transfer, while *A1* is also necessary for host DNA degradation. In contrast, *dmp* and *hdi* (orange) are dispensable for infection. White arrows represent genes whose role in infection is unknown. Black arrows represent three promoters that were experimentally characterized, two overlapping fully or partially with gene *hdi* (P_H 22_ and P_D/E 20_, respectively) and one upstream of gene *011* (12, 13).

The viral genome is delivered in two distinct phases. During the First-Step Transfer (FST), approximately 8% of the genome enters the host cell (Fig. 1) (5). DNA transfer temporarily halts at the “injection stop signal”, a non-coding region rich in repeats and palindromes (6). During this pause, viral pre-early genes are expressed, inhibiting bacterial nucleases and triggering rapid degradation of host DNA (7–10). After 3–5 minutes, DNA transfer resumes in the Second-Step Transfer (SST), delivering the remaining 92% of the genome. The infectious cycle then proceeds with DNA replication, virion assembly and host lysis, culminating ca. 40 minutes post-infection (11).

The FST-DNA harbors 17 predicted open reading frames, named pre-early genes (Fig. 1). Among these, *A1* and *A2*, which encode a putative metallo-phosphatase and a DNA-binding protein, respectively, are essential for infection and SST (7, 14, 15). In contrast, *dmp* (5ʹ-deoxynucleoside monophosphatase) and *hdi* (host division inhibitor) are dispensable for infection (15–17). Ectopic expression of 13 pre-early genes in *E. coli* revealed that *hdi*, gene *011* and gene *015* are toxic in the absence of other viral factors (18). Further analysis showed that *hdi* inhibits host cell division, likely by targeting FtsZ (17), while gene *015* encodes a nuclease that nicks uracil-containing DNA upon its association with the Uracil DNA glycosylase Ung (18).

Beyond its distinctive biphasic DNA delivery, phage T5 is remarkable for its rapid degradation of host DNA: within four minutes of infection, half of the bacterial genome is digested, a process dependent on RNA and protein synthesis and the essential gene *A1* (2, 7, 19). This suggests that viral pre-early genes carried by the FST-DNA play a central role in regulating host DNA degradation. Unlike phages T4 and T7, which recycle nucleotides liberated from the slower host DNA digestion, T5 degrades them into nucleosides via the enzyme Dmp, and, while byproducts are excreted from the cell, it synthesizes nucleotides de novo for its own genome replication (20, 21). Thus, T5 provides a unique system for dissecting host takeover mechanisms and identifying viral effectors that specifically target host functions, including bacterial DNA.

However, critical questions remain: How do T5 pre-early genes influence bacterial physiology and infection dynamics? What are the genetic determinants for the DNase that rapidly degrades the bacterial chromosome? In this study, using ectopic expression and reverse genetics, we identify four additional T5 pre-early genes that are toxic to *E. coli* and pinpoint the primary viral nuclease gene both necessary and sufficient for *in vivo* bacterial genome digestion.

## Results

### Impact of ectopic expression of pre-early genes in *E. coli* on growth

As a first approach in understanding the function of T5 pre-early genes, we systematically studied the impact of their expression on bacterial growth and morphology. The seventeen annotated pre-early genes were individually cloned into plasmid pBAD24 under the control of an arabinose-inducible promoter. Cloning of genes *A1*, *hdi*, *011* and *015* required numerous attempts and the inclusion of glucose in the selective medium, hinting at the potential toxicity of these gene products in the cell. Although the wild-type gene *011* could not be cloned, cloning of a synthetic gene with synonymous codons (but only 76% nucleotide identity, Fig.S1) was successful.

#### Seven T5 pre-early genes are toxic when expressed in *E*. *coli*

*E*. *coli* strain F cells were transformed with pBAD24 derivatives, grown to mid-exponential phase and then serially diluted before spotting on selective solid media containing either glucose or arabinose. Arabinose but not glucose led to reduced bacterial recovery for strains carrying *dmp*, *A1*, *hdi*, *hegG*, *011*, *013* and *015* (Fig. 2).

**Figure 2.**
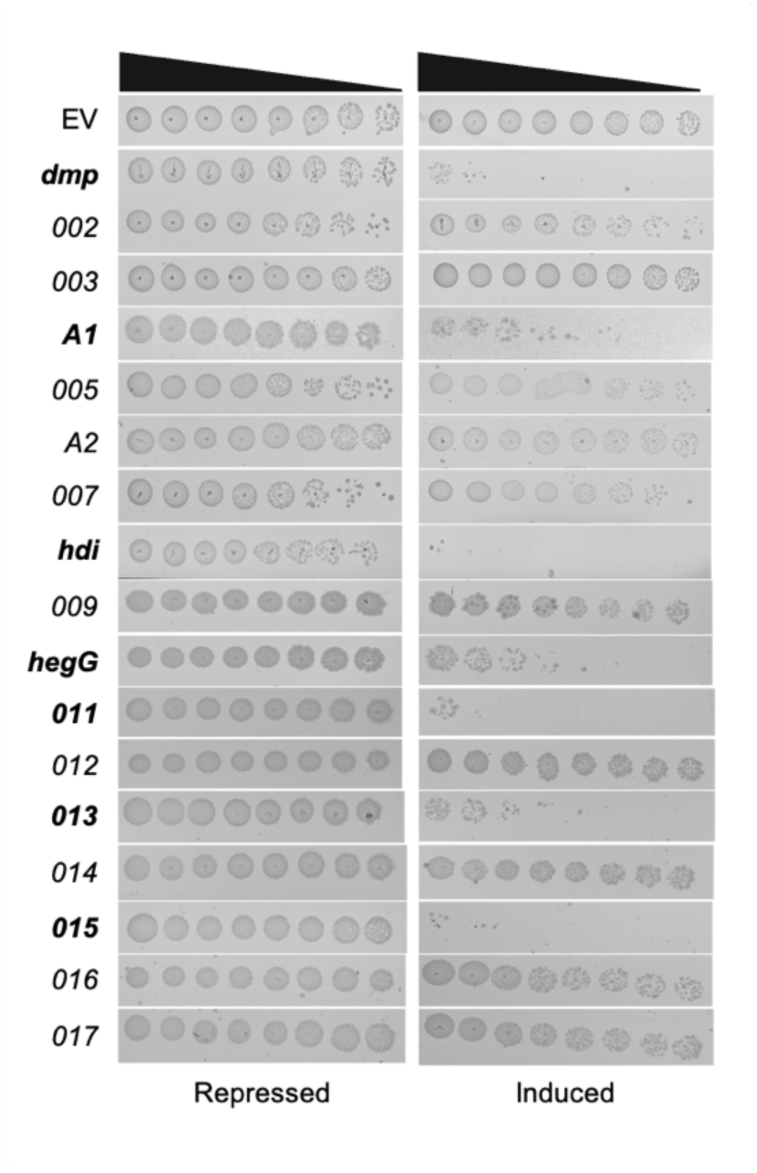
Toxicity of T5 pre-early gene expression in *E. coli* strain. **F.** Bacteria transformed with a pBAD24 vector carrying the indicated pre-early genes were grown overnight at 37 °C in LB medium supplemented with ampicillin and glucose. Overnight cultures were diluted to an OD_600_ of 0.1 in LB with ampicillin and grown for 1 hour at 37 °C. Serial dilutions were then prepared, and 5 µL of each dilution were spotted, from undiluted to most diluted (left to right), onto LB ampicillin agar plates containing either 0.4% D-glucose (gene repression, left panel) or 0.4% L-arabinose (gene induction, right panel). Plates are shown after 24 hours of incubation at 37 °C. EV, empty vector.

Similar results were obtained in another strain of *E*. *coli*, namely DH5α (Fig.S2). Taken together, our results indicate that expression of a subset of T5 pre-early genes is toxic in *E*. *coli* during growth on solid media.

#### Impact of ectopic expression of toxic pre-early genes on bacterial cell morphology, nucleoid organization and DNA content

To investigate the cellular effects of T5 pre-early genes, we examined *E. coli* F cell morphology and nucleoid organization using phase-contrast and fluorescence microscopy of fixed bacteria stained with the DNA-specific dye DAPI. Ectopic expression of these genes induced distinct phenotypic changes, including alterations in cell shape, nucleoid distribution, and DNA content (Fig. 3).

**Figure 3.**
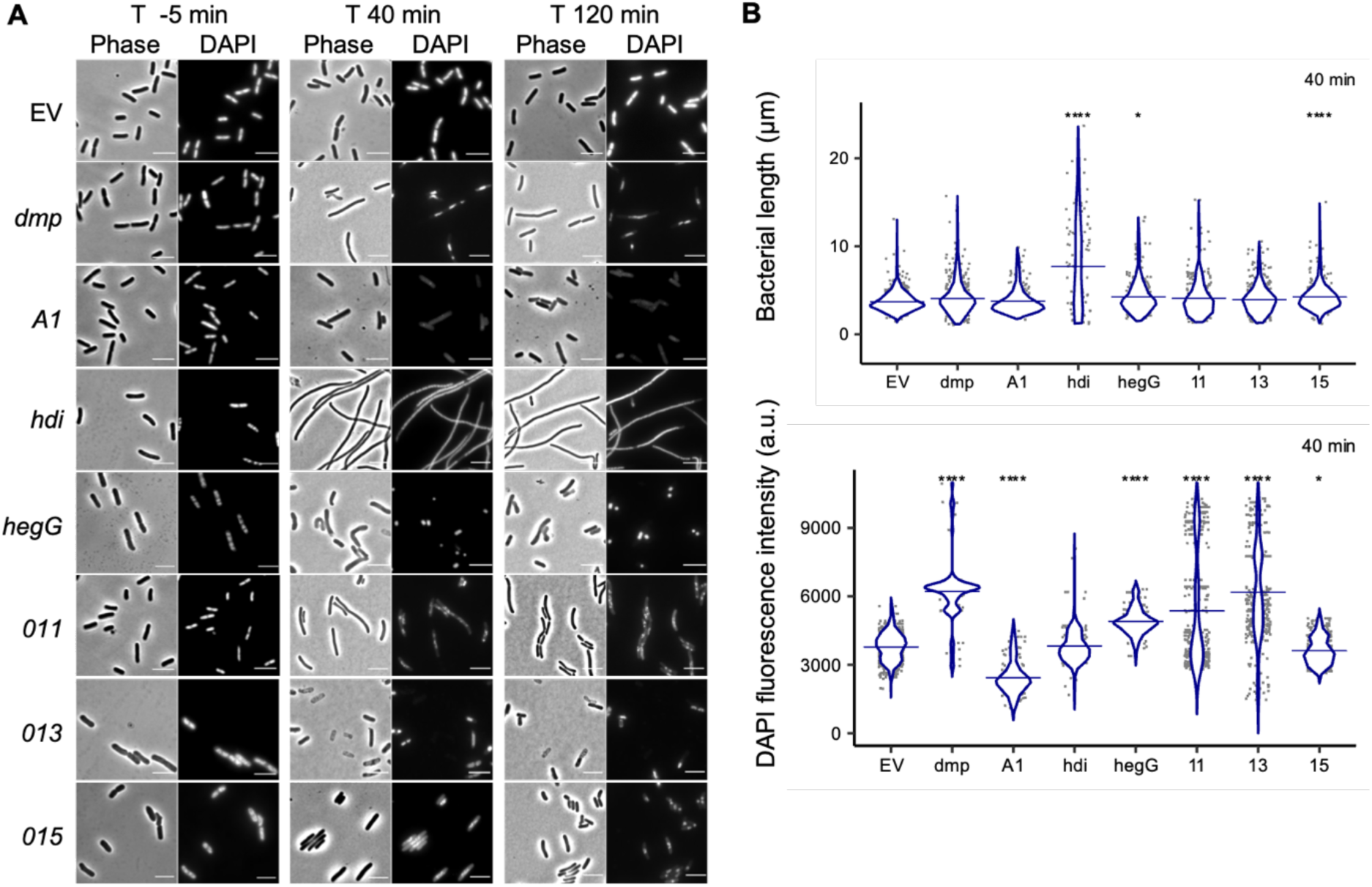
Morphology and DNA staining changes in *E. coli* F cells expressing T5 toxic pre-early genes. **(A)** Phase contrast (Phase) and fluorescence (DAPI) microscopy of *E*. *coli* F cells carrying pBAD24-plasmid derivatives with the indicated viral genes. Cells were fixed and stained with DAPI at 5 minutes pre-induction (-5 min) and at 40 and 120 minutes post-arabinose induction. Time points relative to induction are indicated above each panel. Scale bars, 5 μm. EV, empty vector. **(B)** Violin plots showing bacterial cell length (upper panel) and DAPI fluorescence intensity (lower panel; a.u., arbitrary units) for at least 200 cells after 40 minutes of arabinose induction (n = 2 experiments). Horizontal bars indicate the mean. Statistical significance was determined using a Kruskal-Wallis with Dunn’s post-hoc test, using a Bonferroni correction (*P<0.05; ****P<0.001).

Ectopic expression of *dmp*, which encodes a 5′-deoxynucleoside monophosphatase (22), resulted in cells with bright, diffuse, and centrally located DNA staining (Fig. 3). This phenotype aligns with those reported under thymine starvation or reduced dTTP pools (23), suggesting that *dmp*-mediated cytotoxicity may stem from disrupted nucleotide balance.

In contrast, expression of the predicted metallo-phosphatase gene *A1* caused a dramatic reduction in DAPI staining often confined to a single punctate focus (Fig. 3). These findings suggest that expression of *A1* is sufficient to trigger extensive DNA degradation.

The most pronounced morphological changes were observed upon expression of *hdi*, which induced extensive filamentation with nucleoids distributed along the cell axis (Fig. 3), indicating that *hdi* disrupts normal septation without inhibiting DNA replication. This phenotype mirrors the effects of FtsZ depletion or inhibition (24). Consistent with these observations, expression of *hdi* has been shown to destabilize FtsZ rings during bacterial growth (17).

Expression of *hegG*, a putative HNH endonuclease, led to slightly elongated cells, some of which exhibited pronounced curvature (Fig. 3). Microscopy further showed centrally-located compacted nucleoids with intense DAPI staining, indicative of DNA supercompaction, a hallmark of severe DNA damage (25).

Cells expressing *011* were slightly elongated, with uneven DAPI staining within cells as well as differences between cells, suggesting disrupted nucleoid organization and segregation (Fig. 3).

Expression of *013* produced a heterogeneous population of cells: while some cells remained intact, others appeared shrunken, flattened and devoid of refractive properties, consistent with lysis (Fig. 3). DAPI staining was absent in these lysed cells, whereas intact cells retained DNA. Among the toxic proteins studied here, Gp013 is unique in possessing two predicted transmembrane domains, which might contribute to its lytic activity.

Finally, expression of *015*, an Ung-dependent DNA nickase, resulted in elongated cells, with mid-cell DAPI staining after 40 minutes post-induction (Fig. 3). By 120 minutes, microscopy revealed a mixed population, including cells arrested mid-division with “guillotined” nucleoids trapped at the septum, a phenotype associated with aberrant septation in *E. coli* (26, 27). Other cells displayed nucleoids located toward the poles, consistent with incomplete segregation. Similar elongation and nucleoid condensation were previously reported upon *015* ectopic expression in *E. coli* BW25113 (18), suggesting a conserved mechanism of growth inhibition across strains.

Collectively, these findings revealed the diversity of phenotypic consequences of toxic T5 pre-early gene expression in *E. coli*, including abnormal septation, cell lysis, and disrupted nucleoid organization or segregation. Of particular interest, given our focus on understanding the rapid degradation of bacterial DNA by T5, are three gene products that may play a key role in promoting DNA degradation *in vivo*: the putative metallo-phosphatase A1, the predicted HNH endonuclease HegG, and the Ung-dependent nickase Gp015.

#### Fate of bacterial DNA upon expression of viral nuclease genes *A1*, *hegG* and *015* in *E. coli* strains

To further quantify the dynamics of DNA degradation and cellular changes induced by expression of T5 putative nucleases at the population level, we employed flow cytometry to analyze DNA content and cell size in a high-throughput manner. At different times of gene induction, *E. coli* strain F cells were fixed and stained with propidium iodine (PI). Flow cytometry revealed that the mean PI fluorescence of *A1*-expressing cells dropped by half by 40 minutes of induction, (Fig. 4), suggesting extensive chromosomal degradation.

**Figure 4.**
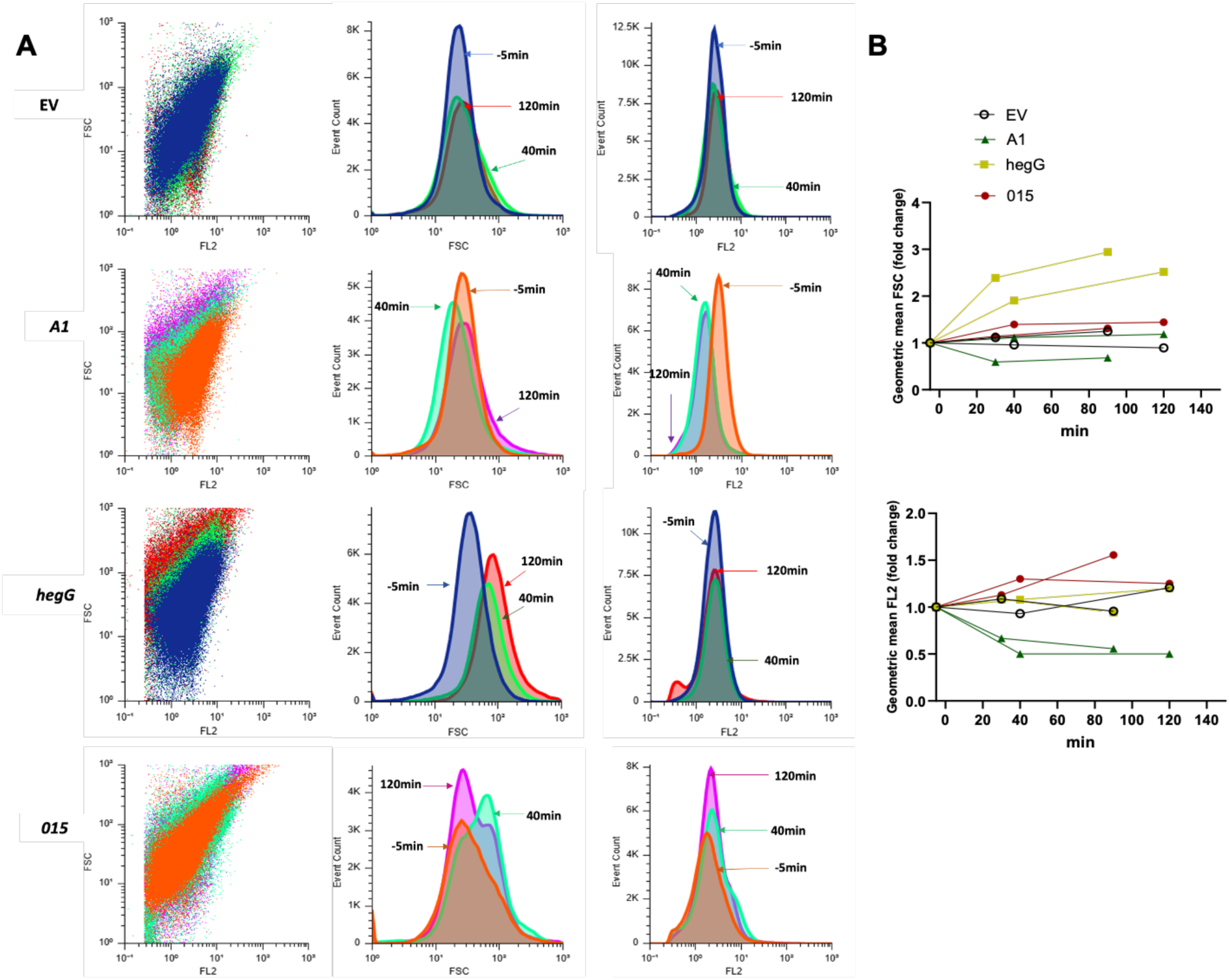
Flow cytometry analysis of *E*. *coli* F cells expressing putative nuclease genes. Cells carried either the empty vector (EV) or one of the putative nuclease genes (*A1*, *hegG*, or *015*). **(A)** Scatter plots and histograms showing DNA content (propidium iodide staining, FL2) and cell size (forward scatter, FSC). Cells were fixed and stained at three time points: 5 minutes before arabinose induction (baseline), and after 40 and 120 minutes post-induction. Each color corresponds to a specific time point. **(B)** Kinetics of cell size (FSC) and DNA content (FL2) from two independent experiments. Data are presented as fold change in geometric mean relative to the baseline (-5 min). The top graph shows changes in cell size, whereas the bottom graph shows changes in PI fluorescence.

Next, we tested the bacterial DNA recovery upon expression of the predicted viral nuclease genes *A1*, *hegG* or *015* in strain DH5α. Total nucleic acids were extracted at different time points, i.e., under non-inducing conditions (5 minutes prior to arabinose addition), and 30 and 90 minutes after induction. Agarose gel electrophoresis revealed the expected pattern of chromosomal and plasmid DNA as well as ribosomal RNA (23S, 16S and 5S) under non-inducing conditions (−5 min) for all transformants (Fig. 5).

**Figure 5.**
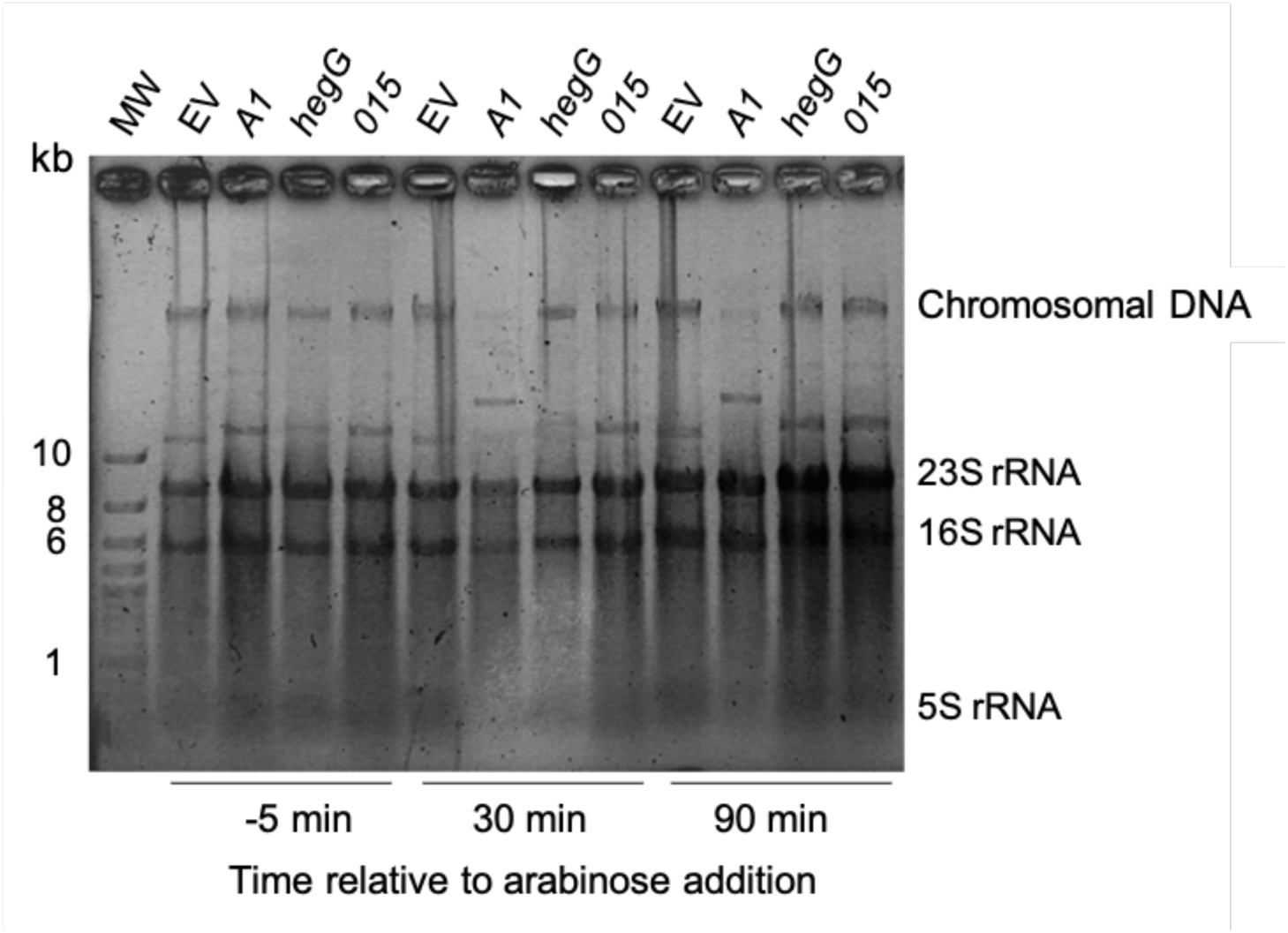
Nucleic acid degradation in *E. coli* DH5α expressing predicted nucleases. Total nucleic acids (DNA and RNA) were extracted at three time points: pre-induction (-5 min), 30 and 90 minutes after induction with 0.2% arabinose. Samples were analyzed by agarose gel electrophoresis and ethidium bromide staining. In *A1*-expressing samples, the band migrating below the chromosomal DNA and above the 10 kb marker was inconsistently observed.

Following induction, marked chromosomal DNA degradation was observed exclusively in *A1*-expressing cells. In contrast, *hegG* and *015* expression did not affect DNA stability (Fig. 5). Recovery of ribosomal RNA was similar in all conditions tested, excluding RNase activity directed against rRNA. Taken together the results suggest that A1 might be a potent nuclease that digests bacterial DNA upon expression and that does not require other viral factors for its activity.

### Construction of T5 mutants and phenotypic analysis

#### At least 12 pre-early genes are dispensable for infection by phage T5

As a next step, we investigated whether other pre-early genes than *A1* may facilitate host DNA degradation during infection. Towards this end, using genome engineering procedures that we recently developed (15), we generated phage mutants and characterized their phenotypes. We started with genes likely located in the same operon than *dmp*, *A1* and *A2* (Fig. 1). To avoid polar effect on downstream genes, deletions were designed to result in open reading frames comprising the first codon and the last ten codons of each mutated gene (Fig. S4). In addition to the deletion mutant in *dmp* already described (15), we could recover mutants deleted in genes *002*, *005*, and *007* (Fig. S3 and S4), indicating that besides the essential genes *A1* and *A2*, four genes downstream of *hdi*, are dispensable for T5 infection. Attempts to delete *hdi* and *009* were prevented by our failure in cloning fragments of this locus that contain the strong promoters P_H 22_ and P_D/E 20_ overlapping fully or partially with *hdi*, respectively (12).

We hypothesized that some pre-early genes might appear nonessential because of putative redundant functions. Such redundancy could be revealed by combining deletions. Hence, we proceeded stepwise to generate phages with multiple mutations. T5 D*257* corresponds to a phage combining the three deletions D*02*, D*05* and D*07*; in parallel, we generated phage T5 DL which carries the 3 deletions D*dmp0203* D*05* D*07* (Fig. S3). Moving towards the second half of the FST-DNA, we also deleted the genes *hegG* to *017* to generate phage T5 DLDR (Fig. S3). This mutant phage lacks most pre-early genes and bears only 4 pre-early genes (*009*, *hdi*, *A2* and *A1*, Fig. S5). Phage DLDR is the only mutant which exhibited smaller plaque size, suggesting that although the 4 remaining pre-early genes are sufficient for infection, some of the missing DNA sequences might facilitate optimal infection. The successful generation of these mutants indicated that thirteen pre-early genes are dispensable for T5 infection.

#### Deletion of pre-early genes alters infection kinetics

Next, one-step growth experiments were conducted to study the infection kinetics of single-gene and multiple-gene mutants in *E*. *coli* strain F (Fig. 6 and S6).

**Figure 6.**
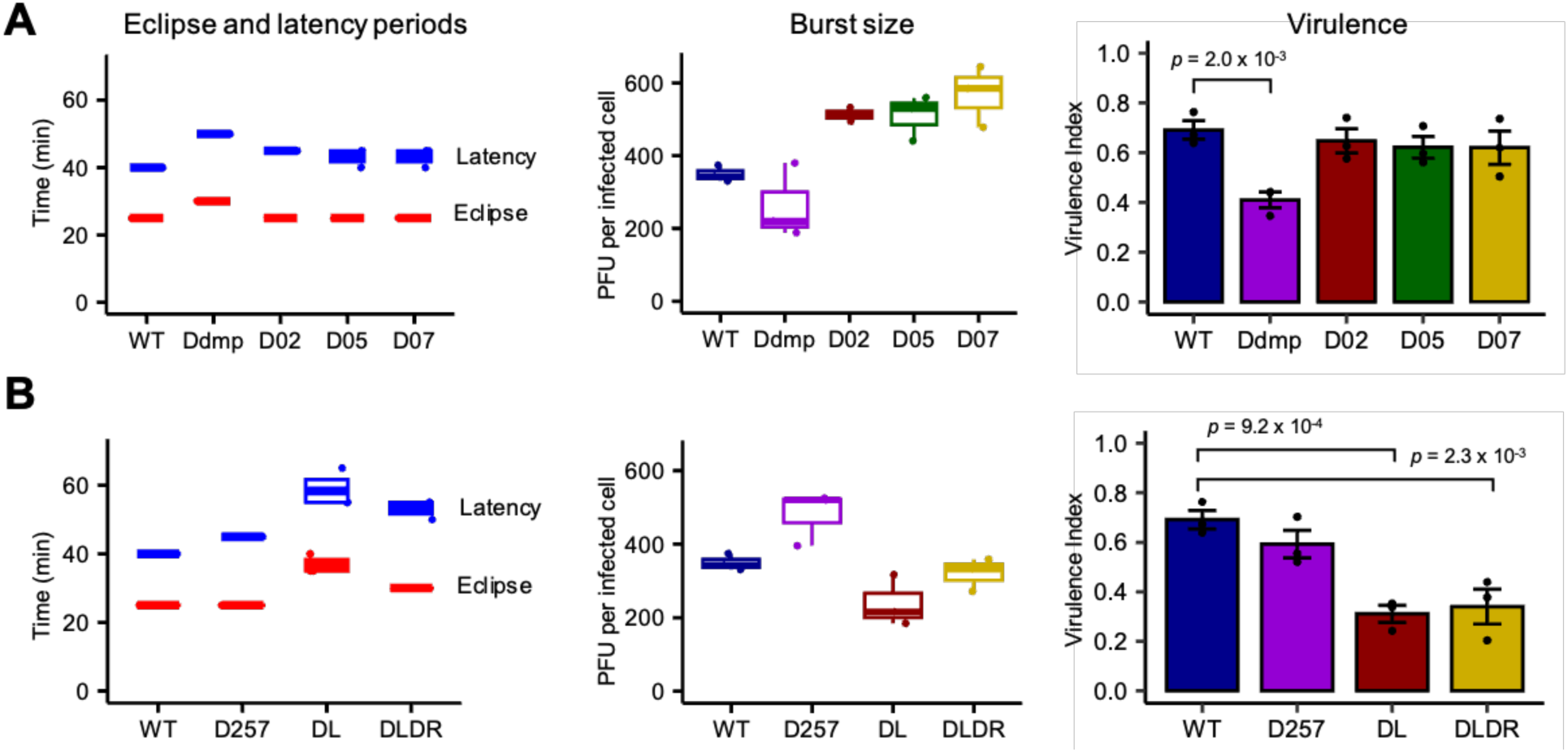
Thirteen pre-early genes are dispensable for infection but their deletion negatively impacts phage virulence. T5 mutants devoid of single **(A)** or multiple **(B)** genes were characterized regarding their eclipse and latency periods (left), burst size (center), or virulence (right) using the *E. coli* strain F. Every parameter was determined thrice and the *p*-values were obtained by ANOVA and Tukey’s Honest Significant Difference (Tukey HSD) post-hoc test. The full comparison of infection timing and burst size is detailed in Table S2. Bacterial reduction curves and local virulence are shown in Fig. S7, while full comparison of virulence index values is listed in Table S3.

Wild-type T5 produced over 350 phages per infection, with an eclipse period of 25 minutes and a latency of 40 minutes, consistent with previous analyses (11, 28, 29). Among mutants carrying single-gene deletions, T5 D*dmp* exhibited the strongest phenotype compared to the wild type, with a small reduction in burst size, and a delay of 5 and 10 minutes in eclipse and latency periods, respectively (Fig. 6 and S6, Table S2). Lack of genes *002*, *005* or *007* was associated with an unaltered eclipse, a slight delay of 5 minutes in latency and a 40 %-increase in burst size relative to wild type (Fig. 6), but only T5 D*02* reached statistical significance for the latter (Table S2).

Taken together, the phenotypes for all the mutants might be classified into 2 categories: (i) mutants with a 5-minute delay in progeny release and a higher burst size (T5 D*02*, T5 D*05*, T5 D*07*, T5 D*257*), (ii) all the mutants devoid of *dmp* (T5 D*dmp*, T5 DL, T5 DLDR), which show longer eclipse and latency periods indicating both virion assembly and progeny release are delayed.

#### Deletion in pre-early genes leads to reduced virulence

To determine the impact of mutations on phage virulence, we followed the optical density of bacterial cultures infected at different multiplicity of infection (MOI) (Fig. S7). These bacterial reduction curves were analyzed to determine the phage virulence index (VI) for the different mutant phages (30). All the mutants devoid of *dmp* (D*dmp*, DL and DLDR) exhibited almost half of the virulence of the wild type (Fig. 6). Conversely, there was no change for the single deletion mutants in *002*, *005* or *007*, or the mutant D*257*, which combines the three deletions (Fig. 6). These data are in agreement with the kinetics of infection described above for these mutants. Taken together, these results indicate that in addition to the essential genes *A1* and *A2*, *dmp* facilitates infection of *E*. *coli* by phage T5.

### Investigating the role of T5 pre-early genes in host DNA degradation during infection

Using the mutant panel described above, as well as the amber mutants in *A1* and *A2* (15), we investigated the role of phage pre-early genes in the degradation of the host DNA by fluorescence microscopy of infected cells. Cells of *E. coli* strain F were infected at an MOI of 10, fixed at different times and stained with DAPI. Infection by the wild-type phage led to a decline in the intensity of the DAPI staining in the first ten minutes, reaching the lowest values at 20 minutes before an increase was observed at 30 minutes, likely due to phage DNA replication. At 30 minutes, the lack of cell refringence indicated that the bacterial integrity was compromised (Fig. 7).

**Figure 7.**
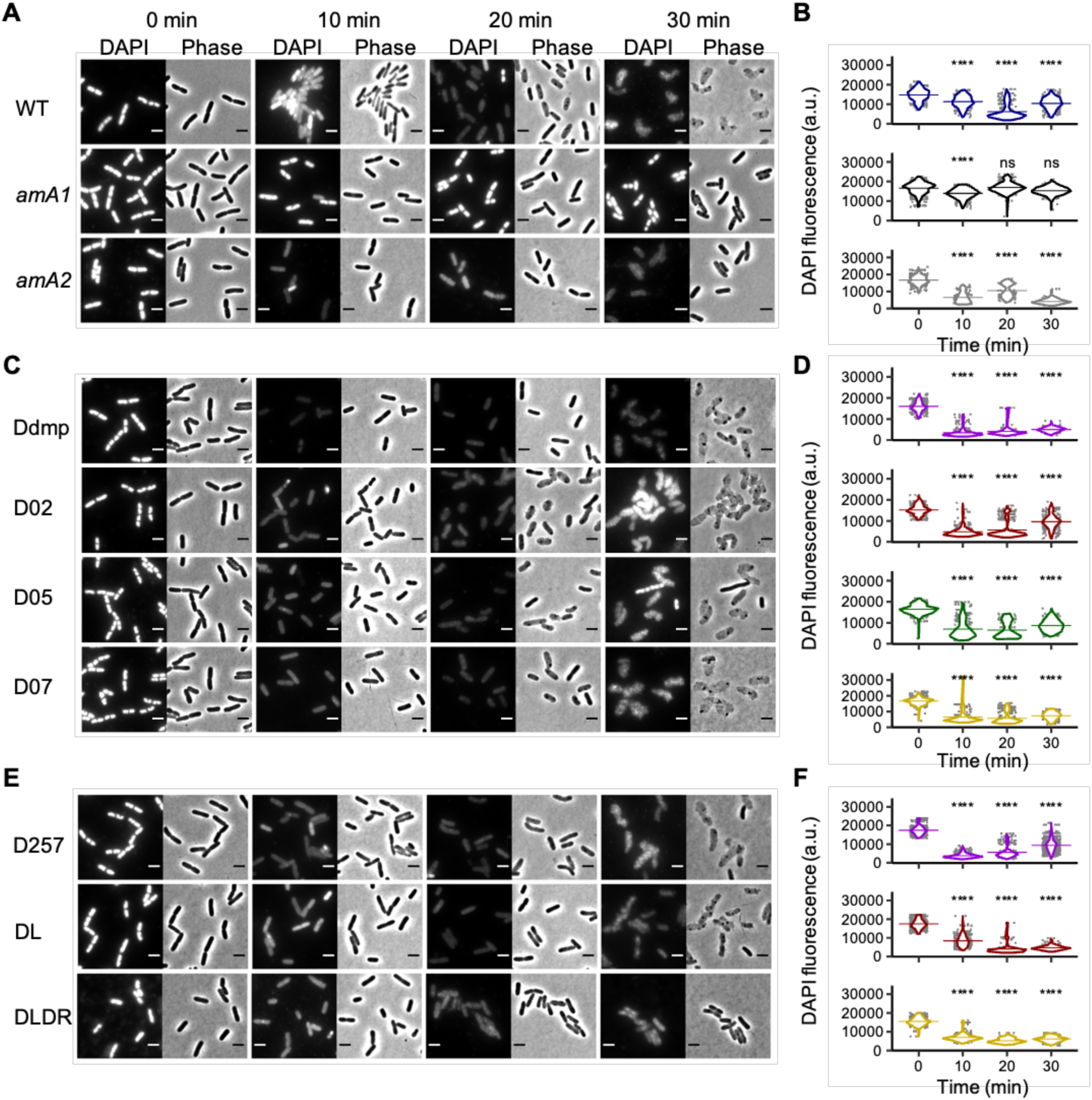
Fluorescence microscopy of *E*. *coli* infected with T5 mutants. Bacterial cultures in exponential phase were infected at MOI of 10, incubated at 37 °C, fixed at different intervals and stained using DAPI. **(A, C, E**) Cells were observed by fluorescence microscopy (DAPI) or using phase contrast (Phase). Scale bar, 2 μm. **(B, D, F)** Violin plots of DNA staining intensity of bacterial cells over time. Results are shown for T5wt and the T5 amber mutants in *A1* and *A2* (panels A and B), single-deletion mutants in pre-early genes *dmp*, *002*, *005* and *007* (panels C and D), and multiple-deletion mutants T5 D257, T5 DL and T5 DLDR (panel E and F). Horizontal bars indicate the mean. Statistical significance was determined using a Kruskal Wallis with Dunn’s post-hoc test (***: *p* < 0.001).

Amber mutants T5 *amA1* and T5 *amA2* led to no lysis after 30 minutes of infection and displayed two distinct phenotypes: DAPI intensity remained roughly constant in cells infected by T5 *amA1*, whereas it rapidly decreased in T5 *amA2*-infected cells. Our observations are consistent with previous reports that *A1* but not *A2* is required to observe host DNA breakdown.

For all the other mutants besides T5 *amA1*, the cellular DNA content decreased to the lowest levels in the first 10 or 20 minutes (Figs. 7C to F) and increased near the end of the infection. Remarkably, the modest rise of DAPI intensity in cells infected by the less-virulent mutants Ddmp, DL and DLDR (Figs. 7D, F), denotes a lower amount of DNA synthesized near the end of the infection, which might explain the delay in the eclipse/latency (Fig. 6).

These observations demonstrate that *A1* is the only pre-early gene required for DNA degradation during infection.

### Distribution of T5 pre-early genes within the *Demerecviridae* family

We investigated then the distribution of the pre-early genes in phage genomes of the genus *Tequintavirus*, the subfamily *Markadamsvirinae* and the family *Demerecviridae*, as it would suggest a conserved strategy for host takeover. Pre-early genes *003*, *009*, *016* and *hegG* were the least frequent, with less than 15% of all *Demerecviridae* genomes analyzed (Fig. 8). While the remaining 13 genes were present in more than 75% of *Tequintavirus* and *Markadamsvirinae* genomes, only seven are found in 60% or more of the *Demerecviridae* genomes, i.e. in decreasing frequency, *A1*, *A2*, *dmp*, *007*, *011*, *005* and *002*. Remarkably, *A1* is present in all 227 *Demerecviridae* genomes analyzed, suggesting it is a hallmark of this family.

**Figure 8.**
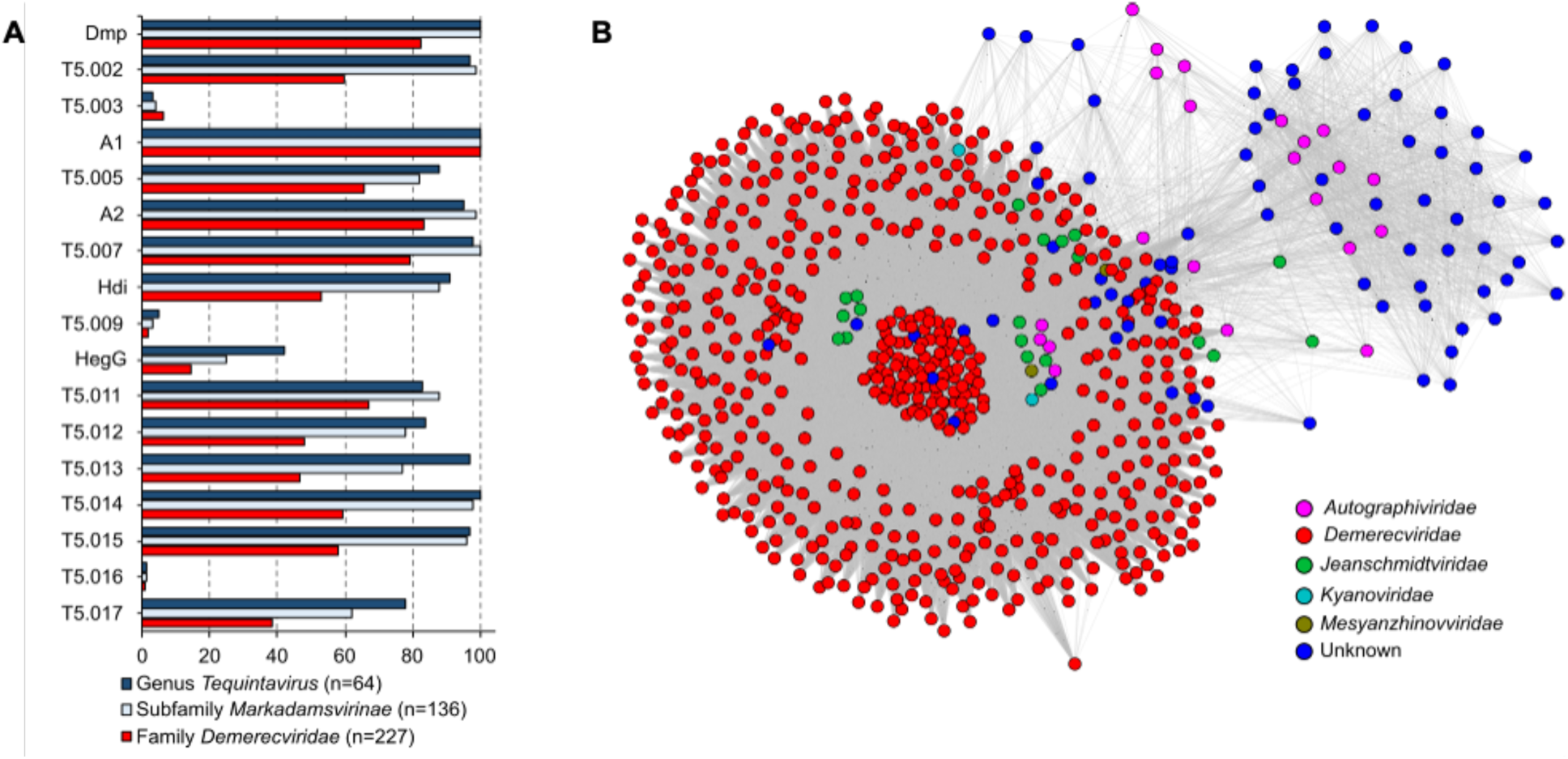
T5 A1 is hallmark of the *Demerecviridae* family. **(A)** Percentage of genomes in T5-related taxa, coding for orthologs to T5 pre-early genes. **(B)** Phage genomes in the INPHARED database coding for proteins homologous to T5 A1. Genomes identified with Blast (tblastn) were then compared with each other with vContACT3.

## Discussion

### Essential and virulence-determining pre-early genes in T5 infection

Our work reveals that, among the fifteen T5 pre-early genes that were analyzed by reverse genetics, only *A1* and *A2* are essential for productive infection. The *dmp* gene, encoding a 5ʹ-deoxynucleoside monophosphatase, enhances phage virulence but is not strictly required. The *dmp* mutant phenotype closely mirrors that described before, where DNA synthesis and lysis were delayed (5, 16). While our phage carried a deletion of most of the *dmp* gene, the molecular basis of the mutation in the previous reports remains unclear. This work and previous studies indicate that Dmp likely facilitates infection by optimizing nucleotide pools, consistent with how its absence reduces the efficiency of host DNA synthesis.

While we successfully constructed T5 mutant phages, including a minimal FST region variant, technical limitations prevented us from generating mutants lacking *hdi* and *009*. However, previous work showed that *hdi* is dispensable for infection (17). Overall, our results, obtained by reverse genetics, align with recent findings from a CRISPR interference assay through RNA targeting, which identified *A1*, *A2*, and *dmp* as important for T5 infection (31). Although the pre-early genes *002*, *003*, *005*, *007*, *hdi*, and *011* to *017* are not required under standard laboratory conditions, we cannot rule out that some of them may become essential under environmental stress or in *E. coli* strains equipped with different defense systems, highlighting the need for further investigation under varied conditions.

### Toxic pre-early genes with roles in cell integrity, septation or nucleoid organization

Among the 13 pre-early genes dispensable for infection, six exhibit toxicity when expressed ectopically in *E. coli*, resulting in phenotypes that can be categorized into three distinct groups. The first category corresponds to cell integrity defects observed upon expression of gene *013*. Given that Gp013 contains two predicted transmembrane domains, we hypothesize that it might disrupt membrane-associated processes, ultimately leading to cell lysis. Notably this toxicity contrasts with the findings of Mahata et al. (2021), who did not report toxicity for gene *013*; differences in experimental conditions, such as expression levels or bacterial strain background, may account for this discrepancy.

A second category encompassed septation defects observed upon expression of *hdi*. Previous work demonstrated that expression of this pre-early gene inhibits cell division by destabilizing FtsZ rings, resulting in filamentous cells (17). This phenotype could be advantageous for T5, as a recent model suggests that induction of filamentation might amplify the replication of phages (32).

A third category comprised changes in nucleoid organization induced by expression of *dmp*, *hegG*, *011*, and *015*. The toxicity of Dmp is likely due to nucleotide pool imbalances, leading to replication defects in *E. coli*. HegG, a predicted HNH endonuclease, triggers DNA supercompaction, a hallmark of severe DNA damage. HNH nucleases are widespread in phage genomes, including T5, which encodes nine such enzymes, although their direct contribution to infection remains unclear. Gp015 functions as a nickase targeting apurinic/apyrimidinic sites in DNA after Ung excises misincorporated uracil bases (18). The final gene in this category, *011*, resists functional predictions and warrants further investigations.

Overall, these observations suggest that toxicity of these non-essential pre-early genes likely reflects their evolved functions in host subversion, either by compromising cellular integrity or rewiring host metabolism to favor phage propagation. Future studies should elucidate how these pre-early proteins fine-tune the T5 infection cycle.

### Essential roles of pre-early genes *A1* and *A2* in T5 phage infection

*A1* and *A2* stand out as the only essential pre-early genes. The A2 protein, a DNA-binding protein, is critical for SST of the phage genome (7, 14). Our work confirmed that it is dispensable for host DNA degradation and demonstrated that its ectopic expression does not impair *E. coli* cell viability. Given its DNA binding properties, A2 may associate with viral DNA and thereby assist in the SST through an as-yet-unknown mechanism.

Among pre-early genes, *A1* emerges as the sole necessary and sufficient factor for host DNA degradation. Predicted to function as a metallo-phosphatase, A1 shares structural features with metal-dependent nucleases involved in double-strand break repair or in the proofreading activity of DNA polymerases. This suggests that A1 could act directly as a DNase mediating the rapid breakdown of the host chromosome. Alternatively, it may activate a host-encoded nuclease. Future experiments will test whether purified A1 protein exhibits nuclease activity in vitro.

### Unique metabolic strategy of T5: host DNA degradation and protection of viral DNA

Notably, A1 exhibits no sequence homology with viral nucleases involved in host DNA digestion by phages T4 or T7 (33, 34) and appears to be predominantly associated with phages belonging to the *Demerecviridae* family. Among characterized phages, T5 employs a distinctive strategy for host DNA degradation, rapidly dismantling the bacterial chromosome within 5 minutes of infection. Unlike phages such as T4 and T7 which repurpose host DNA degradation products for their own replication, T5 excretes these breakdown products (Mozer et al., 1977; Sayers, 2006; Warner et al., 1975), implying a distinct metabolic approach. This observation raises intriguing questions: does T5 deploy host DNA destruction as a “scorched-earth” strategy to suppress the expression of host-encoded defense mechanisms? In this context, while some phages employ nucleases as counter-defense measures, their enzymatic activity is typically sequence-specific (35–37). Given that T5 DNA is not modified (38), it remains unclear how the viral genome evades degradation by A1. This unresolved question underscores the need for further investigation into the molecular mechanisms underlying this protection.

### Pleiotropic functions of A1: regulation of DNA degradation, SST, and transcription

Beyond its role in DNA degradation, A1 also regulates the resumption of viral DNA transfer and interacts with the bacterial RNA polymerase, dampening the transcription from pre-early gene promoters while sparing early gene promoters (13, 39). The pleiotropic functions of A1, including DNA degradation, SST modulation, and potential transcriptional regulation, warrant deeper mechanistic investigation.

Future studies should investigate whether other pre-early nucleases such as Gp015 and HegG, fine-tune the extent or timing of host DNA breakdown and whether their activities are coordinated with A1. Additionally, the environmental and host-dependent essentiality of other pre-early genes merits further examination, as does the molecular interplay between T5 nucleases and host DNA repair systems. Elucidating these mechanisms will not only clarify T5 infection strategy but also provide broader insights into phage-host interactions and the evolution of viral nucleases.

## Materials and Methods

### Bacteria and phage strains used in this study

The bacterial and phage strains used in this study are described in Table 1. Bacteria were grown in LB (10 g tryptone, 5 g yeast extract and 5 g NaCl per liter) supplemented with 1.5% agar for solid media. When appropriate, antibiotics were used at the following concentrations: ampicillin, 200 mg/mL; chloramphenicol, 50 mg/mL; kanamycin, 100 mg/mL. Prior to phage infections, cultures were supplemented with CaCl_2_ (1 mM) and MgCl_2_ (1 mM). T5 wild type (T5wt) and deletion mutants were propagated using *E. coli* strain F. T5 amber mutants were amplified in *E. coli* strain CR63. Plaques were visualized on a double agar layer using molten LB agar (0.5%) supplemented with the indicator bacteria and the phages.

**Table 1.**
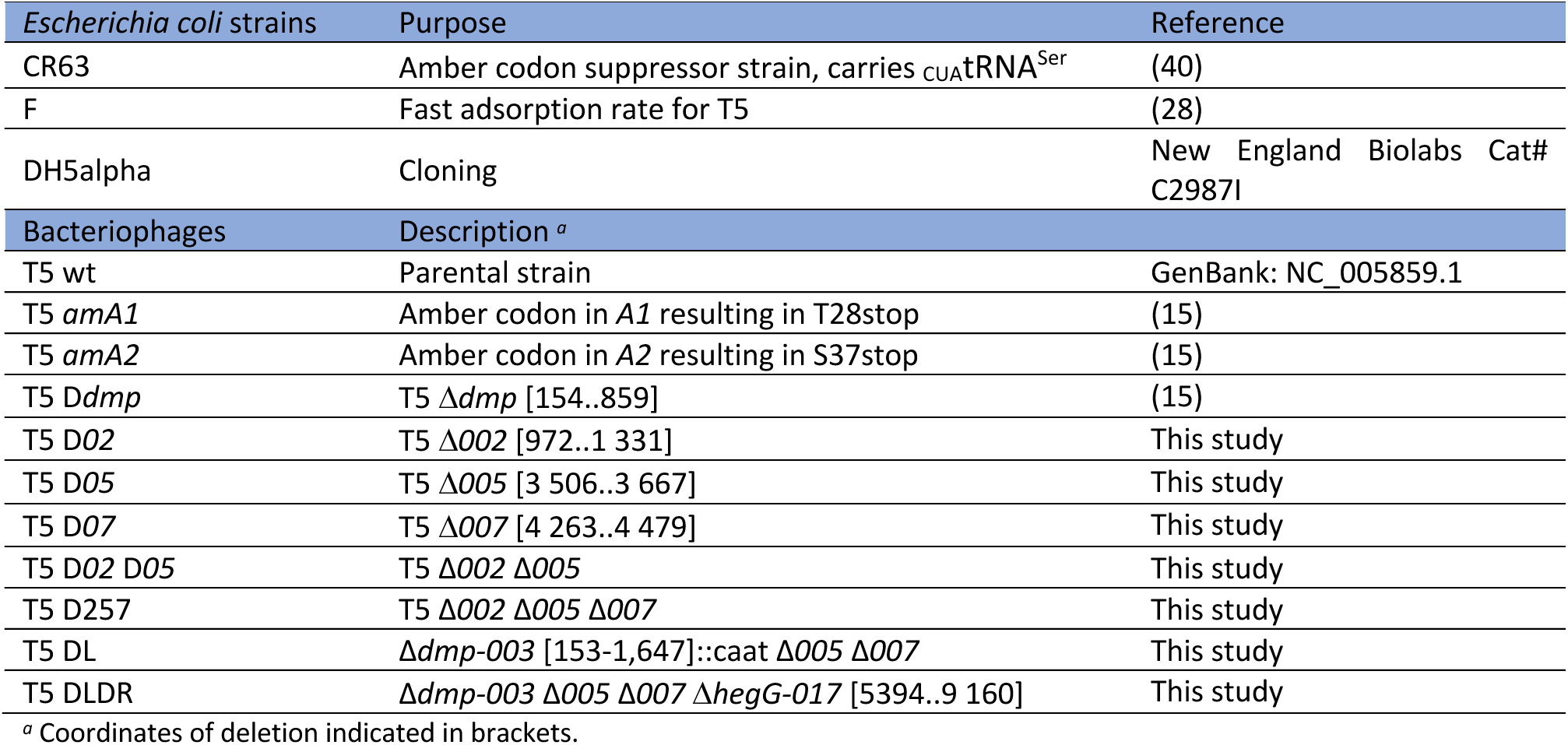
Bacterial and phage strains used in this study.

| *Escherichia coli* strains | Purpose | Reference |
| --- | --- | --- |
| CR63 | Amber codon suppressor strain, carries *CUA*tRNA<sup>Ser</sup> | (40) |
| F | Fast adsorption rate for T5 | (28) |
| DH5alpha | Cloning | New England Biolabs Cat# C2987I |
| Bacteriophages | Description <sup>a</sup> |  |
| T5 wt | Parental strain | GenBank: NC_005859.1 |
| T5 *amA1* | Amber codon in A1 resulting in T28stop | (15) |
| T5 *amA2* | Amber codon in A2 resulting in S37stop | (15) |
| T5 *Ddmp* | T5 $\Delta dmp$ [154..859] | (15) |
| T5 D02 | T5 $\Delta 002$ [972..1 331] | This study |
| T5 D05 | T5 $\Delta 005$ [3 506..3 667] | This study |
| T5 D07 | T5 $\Delta 007$ [4 263..4 479] | This study |
| T5 D02 D05 | T5 $\Delta 002 \Delta 005$ | This study |
| T5 D257 | T5 $\Delta 002 \Delta 005 \Delta 007$ | This study |
| T5 DL | $\Delta dmp-003$ [153-1,647]::*caat* $\Delta 005 \Delta 007$ | This study |
| T5 DLDR | $\Delta dmp-003 \Delta 005 \Delta 007 \Delta hegG-017$ [5394..9 160] | This study |

<sup>a</sup> Coordinates of deletion indicated in brackets.

### DNA designs and clonings

For Golden Gate cloning (41), cyclic ligation-restriction reactions were carried out with 200 ng of plasmid DNA, insert/plasmid ratio 3:1, 1 unit (U) of BsaI and 1 U of T4 ligase, incubated for 30 cycles at 37 °C and 22 °C, 5 minutes each, then 2 hours at 16 °C. An aliquot was transformed in *E. coli* DH5alpha.

To provide viral DNA as a template for homologous recombination during infection, DNA fragments carrying 300-500 bp on either side of the targeted sequence were amplified and cloned into pUCGG (15), all of them with *Bsa*I restriction sites, by Golden Gate cloning. For single gene deletions, to minimize polar effects, we kept the start codon and the last ten codons.

For ectopic expression of the 17 viral genes of the FST-DNA, several strategies were used to clone each individual gene in pBAD24. For the cloning of genes *T5.002*, *T5.005*, *A2* and *T5.007*, DNA fragments were amplified with EcoRI sites at the 5’ ends and XbaI sites at the 3’ ends, digested with EcoRI and XbaI, and then ligated into the pBAD24 vector, which had also been digested with the same enzymes. For the cloning of gene *A1* and *dmp*, similar strategies were employed with sites KpnI/SalI and EcoRI/HindIII, respectively.

To facilitate cloning of the remaining viral genes by the Golden-Gate method in plasmid pBAD24, we replaced the multiple cloning sites with two BsaI sites by PCR amplification with primers pBADFPhos/pBADR and subsequent religation of the amplified fragment, generating plasmid pBAD24-GG. For the remainder viral genes of the FST-DNA, we amplified T5 pre-early genes and cloned them in pBAD24-GG. Primers follow the notation [gene]ATG/[gene]Stop. A codon-optimized version of *T5.011* was synthetized by IDT DNA technologies to be cloned into pBAD24-GG. The plasmids used in this study are listed in Table 2.

**Table 2.** Plasmids used in this study.

| Plasmids |  |  |
| --- | --- | --- |
| pUCGG | used for cloning of DNA template for homologous recombination used in phage genome engineering, ori pMB1(mutant), Amp<sup>R</sup> | (15) |
| pUCDdmp | used for construction of phage T5 *Ddmp* | (15) |
| pUCD02 | used for construction of phage T5 *D02* | This study |
| pUCD05 | used for construction of phage T5 *D05* |  |
| pUCD07 | used for construction of phage T5 *D07* | This study |
| pUCDdmp0203 | used for the construction of phage T5 DL | This study |
| pUCDFST | used for the construction of phage T5 DLDR | This study |
| pAC9 | used in CRISPR-Cas9 interference assay, ori p15A, *araC*, P<sub>BAD</sub>-driven expression of *cas9*, Kan<sup>R</sup> | (15) |
| pAC9-sgRNA02 | pAC9 carrying sgRNA targeting gene *T5.002* | (15) |
| pAC9-sgRNA05 | pAC9 carrying sgRNA targeting gene *T5.005* | (15) |
| pAC9-sgRNA07 | pAC9 carrying sgRNA targeting gene *T5.007* | (15) |
| pBAD24 | used to clone viral genes under control of the P<sub>BAD</sub> promoter, ori pBR322, *araC*, Amp<sup>R</sup> | (42) |
| pBAD24-GG | pBAD24-derived plasmid, compatible with Golden Gate cloning | This study |
| pBAD24-dmp | used for ectopic expression of gene *dmp* | This study |
| pBAD24-002 | ectopic expression of gene *T5.002* | This study |
| pBAD24-003 | ectopic expression of gene *T5.003* | This study |
| pBAD24-A1 | ectopic expression of gene *A1* | This study |
| pBAD24-005 | ectopic expression of gene *T5.005* | This study |
| pBAD24-A2 | ectopic expression of gene *A2* | This study |
| pBAD24-007 | ectopic expression of gene *T5.007* | This study |
| pBAD24-hdi | ectopic expression of gene *hdi* | This study |
| pBAD24-009 | ectopic expression of gene *T5.009* | This study |
| pBAD24-hegG | ectopic expression of gene *hegG* | This study |
| pBAD24-011 | ectopic expression of synthetic gene *T5.011-opt-CDS* | This study |
| pBAD24-012 | ectopic expression of gene *T5.012* | This study |
| pBAD24-013 | ectopic expression of gene *T5.013* | This study |
| pBAD24-014 | ectopic expression of gene *T5.014* | This study |
| pBAD24-015 | ectopic expression of gene *T5.015* | This study |
| pBAD24-016 | ectopic expression of gene *T5.016* | This study |
| pBAD24-017 | ectopic expression of gene *T5.017* | This study |

### Ectopic expression of viral genes in *E. coli*

Bacterial cells transformed with pBAD24-derived plasmids carrying T5 pre-early genes were shaken for 16 hours at 37 °C in LB broth with ampicillin and 0.2% (w/v) D-glucose. These cultures were diluted at OD_600_∼0.1 in LB broth with ampicillin (without glucose), grown for an hour until OD_600_∼0.4 and then further used for fluorescence, flow cytometry or spot assays.

For fluorescence microscopy or flow cytometry assays, 0.2% (w/v) L-arabinose was added to the cultures and aliquots were collected at specified time points.

For spot assays, cells were serially diluted in LB ampicillin. Five μL were then deposited on LB ampicillin agar containing either 0.4% glucose (for gene repression) or 0.4% arabinose (for gene expression) and incubated for 24 hours at 37 °C.

### Fluorescence microscopy

For bacterial cell fixation, 0.5-mL aliquots from cultures were diluted 1:1 with PBS buffer containing 5% formaldehyde and 0.24% glutaraldehyde, incubated at 4 °C for 20 minutes, followed by two washes in phosphate-buffered saline (PBS) and resuspension in 20 μL PBS. A 10 μL aliquot of this suspension was stained with 0.5 μL DAPI (1 mg/mL stock) and mounted on 1.5-mm thick 1% agarose pads prepared in PBS. Fluorescence of DAPI-labelled DNA (excitation at 350 nm, emission at 460 nm) and cell morphology (phase contrast) were visualized using a Zeiss Axio Observer Z1 inverted epifluorescence microscope using an oil immersion 63x NA 1.40 or 100X NA 1.45 objective. Images were acquired using a cooled camera (Hamamatsu Orca Fusion) and the Zeiss ZenBlue 2.3 pro software. Microscopy images were uniformly processed using Fiji/ImageJ, and quantitative analyses (cell length and DAPI intensity for at least 120 cells) were carried out with the plugin MicrobeJ (43). Data were analyzed with R version 4.5.2 and RStudio 2026.01.0.

### Flow cytometry

For fixation of bacterial cells, 1-mL aliquots of bacterial cultures were mixed with 5 mL of ice-cold 70% ethanol and stored at 4 °C for at least 24 hours. Fixed cells were pelleted by centrifugation (10,000 × g, 5 minutes), washed twice with sterile-filtered PBS, and resuspended in 100 μL PBS containing propidium iodide (10 μg/mL) and incubated for 2-3 hours at 37 °C to allow DNA staining.

Flow cytometric acquisition was performed using a CyFlow® Space instrument (Sysmex Partec) equipped with FlowMax software (v2.52). Forward scatter (FSC) was used to estimate cell size, while propidium-iodide fluorescence was excited with a (488-nm) laser and collected in the FL2 channel using a (∼585/542-nm band-pass filter). Signals were recorded on a logarithmic scale, and at least 10,000 events were acquired for each experimental condition.

Flow cytometry data (FCS format) were analyzed using Floreada (https://floreada.io, accessed 2025), a web-based cytometric analysis platform. Debris and doublets were excluded by appropriate gating of FSC parameters, and histograms of FSC and FL2 fluorescence distributions were generated. The geometric mean fluorescence intensity (FL2) and FSC values were calculated and compared between conditions to assess changes in DNA content and cell morphology associated with induced gene expression.

### Phage mutant construction and validation

To generate phage mutants, *E. coli* cells were transformed with template plasmids carrying viral DNA with the desired mutation. Following infection with T5, phage mutant progeny were enriched either through counterselection by CRISPR-Cas9 interference (for deletion of T5.002, T5.005 and T5.007) or through DAS (dilution-amplification-screening) as previously described (15). Verification of the phage mutants was carried out by PCR (Fig. S3) and Sanger sequencing (Fig. S4). Phage mutant stocks were prepared by Cesium chloride gradient as previously described (44).

Whole genome sequence was determined for phage mutants bearing multiple-gene deletions, i.e. T5 DL and T5 DLDR, along with that of the parental strain T5 wt.

To purify phage DNA, 1 mL of CsCl-purified phages (5.10^11^ PFU/mL) was incubated with 40 μL of 0.5 M EDTA, 5 μL of proteinase K (18.7 mg/mL) and 25 μL of 20% SDS at 55 °C for 60 minutes with vigorous vortexing at 20-minute intervals. The mixture was extracted with 1 volume of phenol:chloroform:isoamyl alcohol (PCI, 25:24:1), centrifuged for 5 minutes at room temperature at 10,000 x g. Extraction of the aqueous phase with PCI was repeated. Phage DNA was then precipitated using 2 volumes of ethanol, and sodium acetate was added to a final concentration of 100 mM. Following centrifugation for 10 min at 10,000 x g, the pellet was rinsed with 70% ethanol, air-dried and resuspended in 500 μL H_2_O.

Illumina whole genome sequencing was performed at the NGS platform of the I2BC. Libraries were prepared with TruSeq genomic, NextSeq 500/550 Mid Output Kit v2, with 80 cycles. Between 6-14 million reads were gathered for each sample. Raw data were submitted to the European Nucleotide Archive (ENA) under project PRJEB104473. Samples were T5 wild type (ERS28155442), mutants DL (ERS28155443) and DLDR (ERS28155444).

To map the newly formed sequences as well as to see the deleted regions, a composite sequence of the T5 genome was created: the left terminal repeat was modified according to the results of Sanger sequencing of each mutant and the right terminal repeat was left with no modifications. Short reads were mapped to each reference genome using Bowtie2 in default mode, SAMtools, BEDTools, as described in (45) using a Windows Subsystem Linux (WSL). The results were visualized with the Integrative Genomics Viewer (IGV) version 2.8.9 (Fig. S5).

### One-step growth

The one-step growth assay protocol was adapted from Kropinski (2018) (46). Overnight cultures of the host strain (*E. coli* F) were diluted to OD_600_∼0.1 in LBCaMg (LB supplemented with 1mM CaCl_2_ and 1mM MgCl_2_) and incubated over 1 hour until reaching an OD_600_ of ∼0.4. The cells were infected at an MOI of 10^-3^ for 5 minutes at 37 °C with shaking at 220 rpm and then diluted 10^2^, 10^3^ and 10^4^-fold in prewarmed broth. Diluted cultures were incubated at 37 °C and shaking. Phage titers were determined every 5 minutes using 100 µL aliquots for the double agar layer method. Eclipse was determined in parallel by treating aliquots with chloroform before titration. Assays were carried out thrice: Burst size was calculated as a ratio between the average of three final (highest) timepoint titers over that of the first three (lowest) timepoint titers; eclipse and latent times were determined as the last timepoint preceding a 10-fold titer rise.

### Phage virulence index determination

To determine the phage virulence index (30), phages were diluted to from 10^9^ to 10^2^ PFU/mL in LBCaMg and 100 µL of each dilution was distributed in three wells of a 96-well plate. A pre-culture of *E*. *coli* strain F (OD_600_∼0.4) was adjusted to OD_600_∼0.1 and 100 µL were distributed in each well. The OD_600_ was measured in a Tecan plate reader for 6 hours every 10 minutes. Results were plotted as OD_600_ versus time (h) grouped by phage, to visualize the bacterial reduction curves (Fig. S7A). The area under the bacterial reduction curves (AUC) at each MOI were normalized to the AUC of the uninfected culture to obtain the local virulence as a function of the MOI. Finally, for each phage tested, its virulence index was calculated as the area under the local virulence curve over the range of MOI. Area under the curves were estimated using the R function MESS::auc. R version R 4.5.2.

### Comparative genomics

We identified genomes encoding for proteins homologous to A1 within the *Demerecviridae* family (taxid:2731690) and beyond using Blast (tblastn) against the NCBI core_nt database, with default parameters. We selected the identified genomes from the INPHARED database (version 14Apr2025; https://github.com/RyanCook94/inphared) and employed vConTACT3 (https://bitbucket.org/MAVERICLab/vcontact3/src/master/) for genome clustering based on the shared gene content. The exported data were visualized as a cosmograph using ggraph package in R version 4.5.2, using Large Graph Layout (LGL). Final rendering was performed with Inkscape version 1.0.1.

## Recognition

We acknowledge Madalena Renoir for assistance with cloning. We acknowledge the sequencing and bioinformatics expertise of the I2BC High-throughput sequencing facility, supported by France Génomique (funded by the French National Program “Investissement d’Avenir” ANR-10-INBS-09).

## Authors contributions

**Godfred Morkeyile Annor -** Conceptualization, Investigation, Visualization, Writing-review and editing

**Luis Ramirez-Chamorro -** Conceptualization, Formal analysis, Investigation, Visualization, Writing-review and editing

**Leo Zangelmi** - Investigation

**Pascale Boulanger** - Conceptualization, Supervision, Funding acquisition, Writing-review and editing

**Ombeline Rossier -** Conceptualization, Supervision, Writing-original draft, Writing-review and editing

