## Supplementary data for "Dissecting the genetic determinants of bacterial DNA degradation by bacteriophage T5"

Table S1. Oligonucleotide primers used in this study.
Table S2. Pairwise comparison[a] of one-step growth parameters using ANOVA and Tukey's Honest Significant Difference post-hoc test
Table S3. Dunn's post-hoc test for comparison of phage virulence [a]

Figure S1. Cloned synthetic construct encoding gene product Gp011.
Figure S2. Toxicity assay of T5 pre-early genes in *E. coli* strain DH5α.
Figure S3. PCR amplification of targeted locus for the different deletion mutants.
Figure S4. Sanger sequencing from mutant strains showing the deletion sites in *dmp*, *002*, *005*, *007* and from *hegG* to *017*.
Figure S5. Verification of mutants DL and DLDR by whole genome sequencing (WGS).
Figure S6. One-step growth curves from mutants of T5.
Figure S7. Bacterial reduction and local virulence curves.

**Table S1.** Oligonucleotide primers used in this study.

| Homologous recombination | | Sequence[a] |
|---|---|---|
| Deletion - *dmp* | dmpF | aaaaggtctcaaaaaGGCGGGGTGGATTTAATGGA |
| | dmpR | aaaaggtctcaccccCTGGCGTGACTGGTAGTCTG |
| | DdmpF | aaaaggtctcacattTTGTTACGTTCTCCATTTGAGG |
| | DdmpR | aaaaggtctcaaatgCAATATATTGAGAAATTAAAAGTTGCGTAATAATTAAAG |
| Deletion - *T5.002* | FC533/52-LR | atgaccatgattacgccCTATCACCACGACCGCGCAA |
| | FC1965/46-LR | gtaaaacgacggccagtTTGAATGCGGCGTACACGGT |
| | D02F | aaaggtctcaTTTATCGCTGATGCGCTAGCGTAT |
| | D02R | aaaggtctcaTAAACATGTTTATAACTCCAATTAGTTTAATAAG |
| Deletion - *T5.005* | 05F | aaaaggtctcaaaaaACCGCAAAATTCGCTTGGAA |
| | 05R | aaaaggtctcaccccCTAACCAGCAAATCAGCGCC |
| | D05F | aaaggtctcacaccCATTTTAAAACTCCAGTCAAAGGG |
| | D05R | aaaggtctcaGGTGTTAGTTCTAGCCTTATGCCTTT |
| Deletion - *T5.007* | 07F-3 | aaaaggtctcaccccTTCCAAGCGAATTTTGCGGT |
| | 07R-3 | aaaaggtctcaaaaaCCCAGTTCGATGAGAGCGAT |
| | D07F | aaaggtctcaCAGCGTAACGAAAAATGGGAAGC |
| | D07R | aaaggtctcagctgCATAATATAAACTCCAGTTTATTAAGGGG |
| *Deletion - dmp* to *T5.003* | 03R | aaaaggtctcaccccATGGTGACACCTACGGGCTTGT |
| | D03F | aaaaggtctcacattCATTTAATTTAGCTCCTGTAATAGCGGC |
| | Ddmp0203-Scr | GCCGCTATTACAGGAGCTAAATTAAATGaatg |
| *Deletion - hegG* to *T5.017* | FST-F | aaaaggtctcaaaaaTACCCAAAAATGGCGCAACC |
| | FST-R | aaaaggtctcaccccGCACGATTCGCCTTGACAAA |
| | DFST-F | aaaaggtctcacgagGATGGTTTAATCACTGGTATAAACGG |
| | DFST-R | aaaaggtctcaCTCGTTTAGTCTTGATTTTTAAAAGTCAATATC |
| | DFSTScr | GTTTATACCAGTGATTAAACCATCctcg |
| Amplification pBAD24 | pBAD24Phos<br>pBADR | [Phos]-aaaaggtctcaTTTGGCGGATGAGAGAAGATTTTC<br>aaaaggtctcaGTGAATTCCTCCTGCTAGCC |
| Cloning into pBAD24-GG | | |
| *dmp* | dmpATG | aaaaaa*gaattc*aacATGAATCAAGTTAAAACGAATATTACCCGT |
| | dmpStop | aaaaaa*aagctt*TTACGCAACTTTTAATTTCTCAATATATTGTTG |
| *T5.002* | ORF02_EcoRI_F | gagttata*GAATTC*aacATGGCTAAATCTAAC |
| | ORF02_XbaI_R | attaacta*TCTAGA*TTAAAGGCCATACGCTAG |
| *T5.003* | 03ATG | aaaaggtctcatcacATGGCTATTAAAATTAATCTTCCCAGCAT |
| | 03Stop | aaaaggtctcacaaaTTATTTGCTAGCTTCAAGACTTACAACAAA |
| *A1* | Oligo pBAD A1F | t*GGTACC*cggggatcctcATGGTTATTTCCGC |
| | Oligo pBAD A1R | gcctgca*GGTCGAC*tcTTACGCAAAGAATTTACC |
| *T5.005* | ORF05_EcoRI_F | ggagttAt*GAATTC*aaaATGATCTTTTACCC |
| | ORF05_XbaI_R | gcaaca*TCTAGA*TTAATTTAAAGGCATAAG |
| *A2* | pBAD A2 EcoRI F | gttac*GAATTC*atcATGACTAACGCTAAAACCG |
| | pBAD A2 XbaI R | taaa*TCTAGA*TTATTCGCCTGCGGCTTG |
| *T5.007* | ORF07_EcoRI_F | gtttat*GAATTC*attATGCAAAACGTTAC |
| | ORF07_XbaI_R | cacta*TCTAGA*TTACGCTACGGCTTC |
| *T5.008* | 08ATG | aaaaggtctcatcacATGAAACAAACTTTATTGATCACTGGTAAAC |
| | 08Stop | aaaaggtctcacaaaTCAAACTATTTTTTGCAGTTTTTTATGATTTTC |
| *T5.009* | 09ATG | aaaaggtctcatcacATGACGAACTACACGGCGG |
| | 09Stop | aaaaggtctcacaaaTCATATCGTCGATTCCTTATAATATATAAG |
| *hegG* | hegGATG | aaaaggtctcatcacATGGGTCTATTTAAAGATATAGATTATTCGC |
| | hegGStop | aaaaggtctcacaaaTTAATTGACAATTCCAAATTCAGCTTTTTTAG |
| *T5.011* | 11ATG | aaaaggtctcatcacATGCGGCTATATAAACCAGATAACG |
| | 11Stop | aaaaggtctcacaaaTTACTCGCTTTCGGTGTTAACGG |
| *T5.012* | 12ATG | aaaaggtctcatcacATGGTTGCTTATTCTTCTTCTGACG |
| | 12Stop | aaaaggtctcacaaaTTACATTAAAACAATTTTGGCGGCTTGTT |

| | | |
|---|---|---|
| *T5.013* | 13ATG | aaaaggtctcatcacATGGTTATTTTTCGAGCTATTCTGGC |
| | 13Stop | aaaaggtctcacaaaTTATTCTCCTTTATTCATGCAGCCC |
| *T5.014* | 14ATG | aaaaggtctcatcacATGATCCGCAACGTTTCTCTTGC |
| | 14Stop | aaaaggtctcacaaaTTATAACACTCCACGAAAAGCCG |
| *T5.015* | 15ATG | aaaaggtctcatcacATGGTAATTTATGAAGGCAATCGCTTTG |
| | 15Stop | aaaaggtctcacaaaTCATTGCGCCACCTTTCTGAAAG |
| *T5.016* | 16ATG | aaaaggtctcatcacATGATTCCGCTGGTGGCGATA |
| | 16Stop | aaaaggtctcacaaaTTAGTTGACTATCTCAAAGATAGCGTTC |
| *T5.017* | 17ATG | aaaaggtctcatcacATGAGCAATAAAATTATTGTGACCAAAACTAC |
| | 17Stop | aaaaggtctcacaaaTCATAAATCCCCCTCGGGTAATC |

[a] Upper-case letters match with the T5 genome or pBAD24 plasmid.

**A** Synthetic DNA fragment for 011-opt-CDS cloning

```
aaaaggtctcatcacATGCGTTTATATAAGCCCGACAACGCAACGGTCCTGAAAGGGGCGCTTCGCAATCTTCTGGAC
GGAAGTCGCACCACATCGATTAAGCATTTCATCAACAAAGCTGAAACCATTCACCAGAATTTTTTCAATGACTATGAT
GCTTACGACATGGATTCATTCTTACCACTGTATAAACAAGGAGCAACTATCATCCTTTACAACCGCGACTGTCACATT
TTCACGGGACGCGGGGCCGGGCCAGATACAGACTTTATTGTTGCCCAAGGTAACAATATTGTCGCATATCACAGCAAT
TCTGGTCGCCAAAGTTTTAATATTAATAACTTCGAACTGGTAGGTTTAGAACTTATTAACGGTTTGAGTAGTTTGGAG
GGCTTCTTAGATGTTATCTGGAAATACTTATCAGCCGGCTGGCACCGCCCAGACTCTGACGGGTCTCAATATACCGCG
GTCATGGCATTTGACGGAGATGGAAACTTTTTGTTGCCGTCCGAACTGTATCCCTATGTGTACAATGAACTTGTGCAA
GAGGCTGGCTTTATCTTTTGGAAAGATGAGAACCAGATCATCGATAACACGGAGAAGCACGCTGTAAACAGTAATACA
CAGAGCGTTAACACTCAAATTACCGAGAACAAGGGGAAAAATGCTATGACAAAGATCGCTAACATTGTAGCCGCCAAT
AAATCGGCTGTAGTTAACGCAGCGAAACTTGAGGCGGGTAAGATTGCACTTACACAAATCACTAAAGTAGCGGCGAAG
AAGGCTCCATTCATGATTAAGGGGTACATCGATACACCGATTGGCCGCGTTGTAATCGCAAACCTGTTATCTGTCGCG
GTCGACCAGTACGCCCCCTCCAACCAAAAGGCCAAAGCAGTCGCCGGTGCGGCAATGGAAGCTGCCATGTTGGAGATG
GTGCAGTCTTTCAACATTGCGGAAATGATTGACGAGATGGTTAAGGGAATCGATATCAGCACGTTTACGGTAAACACA
GAATCAGAATAAtttgtgagaccttttt
```

**B** CLUSTALW 011-opt-CDS

```
011-CDS        atgcggctatataaaccagataacgcaaccgtcttaaagggcgccttgcgcaacctgctg 60
011-opt-CDS    atgcgtttatataagcccgacaacgcaacggtcctgaaaggggcgcttcgcaatcttctg 60
               *****  ******* ** ** ******** *** * ** ** **  * ***** ** ***

011-CDS        gatggtagcaggacaaccagcattaaacatttattaacaaggctgaaactattcaccaa 120
011-opt-CDS    gacggaagtcgcaccacatcgattaagcatttcatcaacaaagctgaaaccattcaccag 120
               ** ** **  * ** **    ***** ***** ** ***** ******** ********

011-CDS        aactttttaatgattatgatgcctatgatatggatagttttctacccctatataagcag 180
011-opt-CDS    aattttttcaatgactatgatgcttacgacatggattcattcttaccactgtataaacaa 180
               ** ***** ***** ******** ** ** ******   **   **** ** ***** **

011-CDS        ggggcaacgattatcttatataatcgtgattgccatatctttacgggcaggggagcgggg 240
011-opt-CDS    ggagcaactatcatcctttacaaccgcgactgtcacattttcacgggacgcggggccggg 240
               ** ***** ** *** * ** ** ** ** ** ** ** ** *****  * ** ** ***

011-CDS        ccggatactgattttattgttgcgcagggtaataatatcgttgcgtatcatagcaattca 300
011-opt-CDS    ccagatacagactttattgttgcccaaggtaacaatattgtcgcatatcacagcaattct 300
               ** ***** ** *********** ** ***** ***** ** ** ***** ********

011-CDS        ggccgtcaatcatttaatataaataattttgagcttgttgggcttgaattaataaacgga 360
011-opt-CDS    ggtcgccaaagttttaatattaataacttcgaactggtaggtttagaacttattaacggt 360
               ** ** ***   ******** ***** ** ** ** ** **   * *** * ** *****

011-CDS        ctttctagccttgagggatttttagatgttatctggaagtatttatctgctggatggcat 420
011-opt-CDS    ttgagtagtttggagggcttcttagatgttatctggaaatacttatcagccggctggcac 420
                *   ***  * ***** ** **************** ** ***** ** ** *****

011-CDS        agacctgatagtgatggatctcaatatacggctgtaatggcttttgatggcgacggtaat 480
011-opt-CDS    cgcccagactctgacgggtctcaatataccgcggtcatggcatttgacggagatggaaac 480
                * ** **   *** ** *********** ** ** ***** ***** ** ** ** **

011-CDS        tttttattaccatcggaattatatccgtatgtatataatgagctagtacaagaagcgggt 540
011-opt-CDS    tttttgttgccgtccgaactgtatccctatgtgtacaatgaacttgtgcaagaggctggc 540
               ***** ** ** ** *** * ***** ***** ** ***** ** ** ***** ** **

011-CDS        tttattttctggaaagatgagaatcagatcattgacaacacggaaaaacatgctgtaaat 600
011-opt-CDS    tttatcttttggaaagatgagaaccagatcatcgataacacggagaagcacgctgtaaac 600
               ***** ** ************* ******** ** ******** ** ** ********

011-CDS        tctaacactcaaagcgttaacactcaaatcactgaaaacaaaggaaaaaatgccatgacc 660
011-opt-CDS    agtaatacacagagcgttaacactcaaattaccgagaacaaggggaaaaatgctatgaca 660
                 *** ** ** **************** ** ** ***** ** ******** *****

011-CDS        aagatcgctaatatcgttgccgctaataaatccgccgttgtaaatgctgcaaaactggaa 720
011-opt-CDS    aagatcgctaacattgtagccgccaataaatcggctgtagttaacgcagcgaaacttgag 720
               ********** ** ** ***** ******** ** ** ** ** ** ** ***** **

011-CDS        gcgggcaaaattgcattaactcaaatcacgaaagtagcggctaaaaaagcaccatttatg 780
```

```
011-opt-CDS   gcgggtaagattgcacttacacaaatcactaaagtagcggcgaagaaggctccattcatg 780
              ***** ** ****** * ** ******** *********** ** ** ** ***** ***

011-CDS       attaaaggttatattgatacgcctattggtcgggtagttattgctaacctgctgagcgta 840
011-opt-CDS   attaagggtacatcgatacaccgattggccgcgttgtaatcgcaaacctgttatctgtc 840
              ***** ** ** ** ***** ** ***** ** ** ** ** ** ****** *    **

011-CDS       gcggttgaccagtacgcccctagcaaccaaaaggcgaaagcggtagcgggtgcagctatg 900
011-opt-CDS   gcggtcgaccagtacgccccctccaaccaaaaggccaaagcagtcgccggtgcggcaatg 900
              ***** **************   ************ ***** ** ** ***** ** ***

011-CDS       gaagcggctatgctagaaatggtgcaaagttttaacatcgctgaaatgattgatgaaatg 960
011-opt-CDS   gaagctgccatgttggagatggtgcagtctttcaacattgcggaaatgattgacgagatg 960
              ***** ** *** * ** ********   *** ***** ** *********** ** ***

011-CDS       gtgaaaggtattgatatttctacttttaccgttaacaccgaaagcgagtaa 1011
011-opt-CDS   gttaagggaatcgatatcagcacgtttacggtaaacacagaatcagaataa 1011
              ** ** ** ** *****    ** ***** ** ***** ***   ** ***
```

**Figure S8. Cloned synthetic construct encoding gene product Gp011.**

**(A)** Sequence used for cloning into pBAD4-GG. The open reading frame appears in capital letters. The underlined sequences were used to facilitate Golden Gate cloning. **(B)** Alignment of wild-type gene *011* coding sequence (011-CDS based on Reference sequence NC_005859.1) with synthetic coding sequence (011-opt-CDS). Sequences exhibit 76 % identity marked with a star and 242 mismatches. They encode an identical protein.

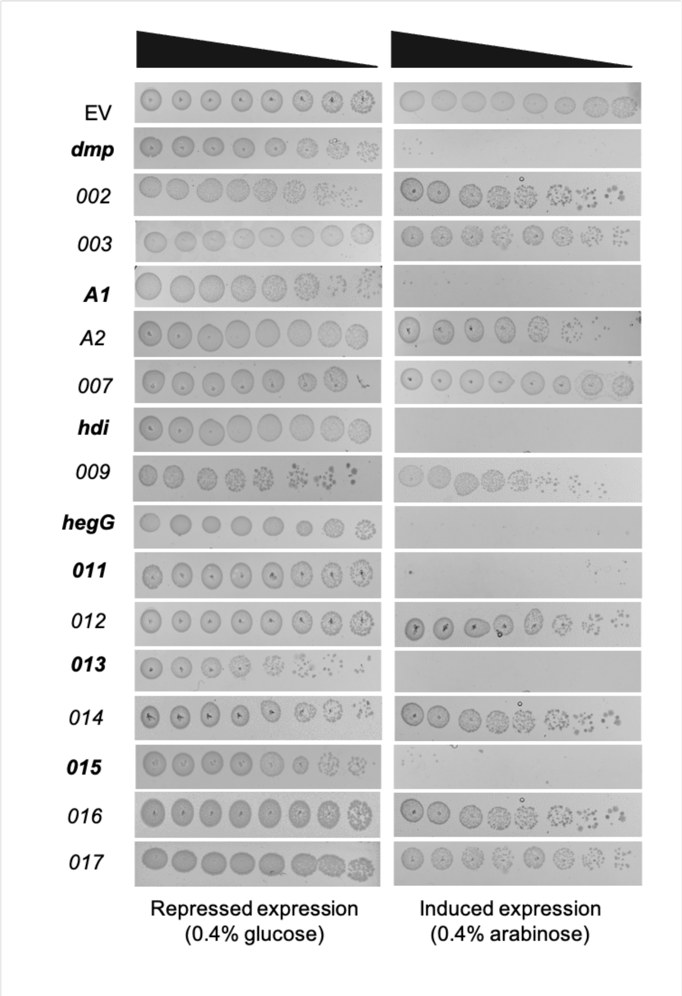


**Figure S9. Toxicity assay of T5 pre-early genes in *E. coli* strain DH5α.**
Bacteria were transformed with pBAD24 derivatives carrying the indicated genes and grown overnight at 37 °C in LB supplemented with ampicillin and glucose. After dilution to $OD_{600}$ 0.1 in LB supplemented with ampicillin, bacteria were grown for an hour at 37 °C, and then serially diluted in LB ampicillin. Five µL were then spotted on LB ampicillin agar containing either glucose (for gene repression, left panel) or arabinose (for gene expression, right panel) and incubated for 24 hours at 37 °C. EV, empty vector.

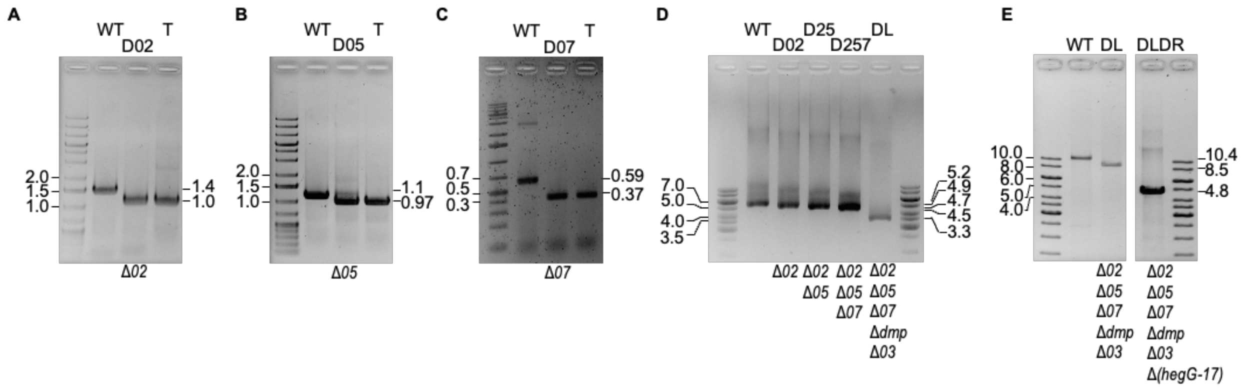


**Figure S10. PCR amplification of targeted locus for the different deletion mutants.**
(A) T5 D*02*, (B) T5 D*05*, (C) T5 D*07*, (D) T5 D257 and DL (T5 D*dmp0203* D*05* D*07*), (E) T5 DLDR (T5 D*dmp0203* D*05* D*07* D*hegG-17*). Amplicon observed with the wild-type phage (WT), the mutant phage and the template plasmid (T). Molecular sizes (kb) from ladders are shown on the left of each gel whereas expected amplicon lengths are shown on the right. Resulting genotype is indicated below the gels.

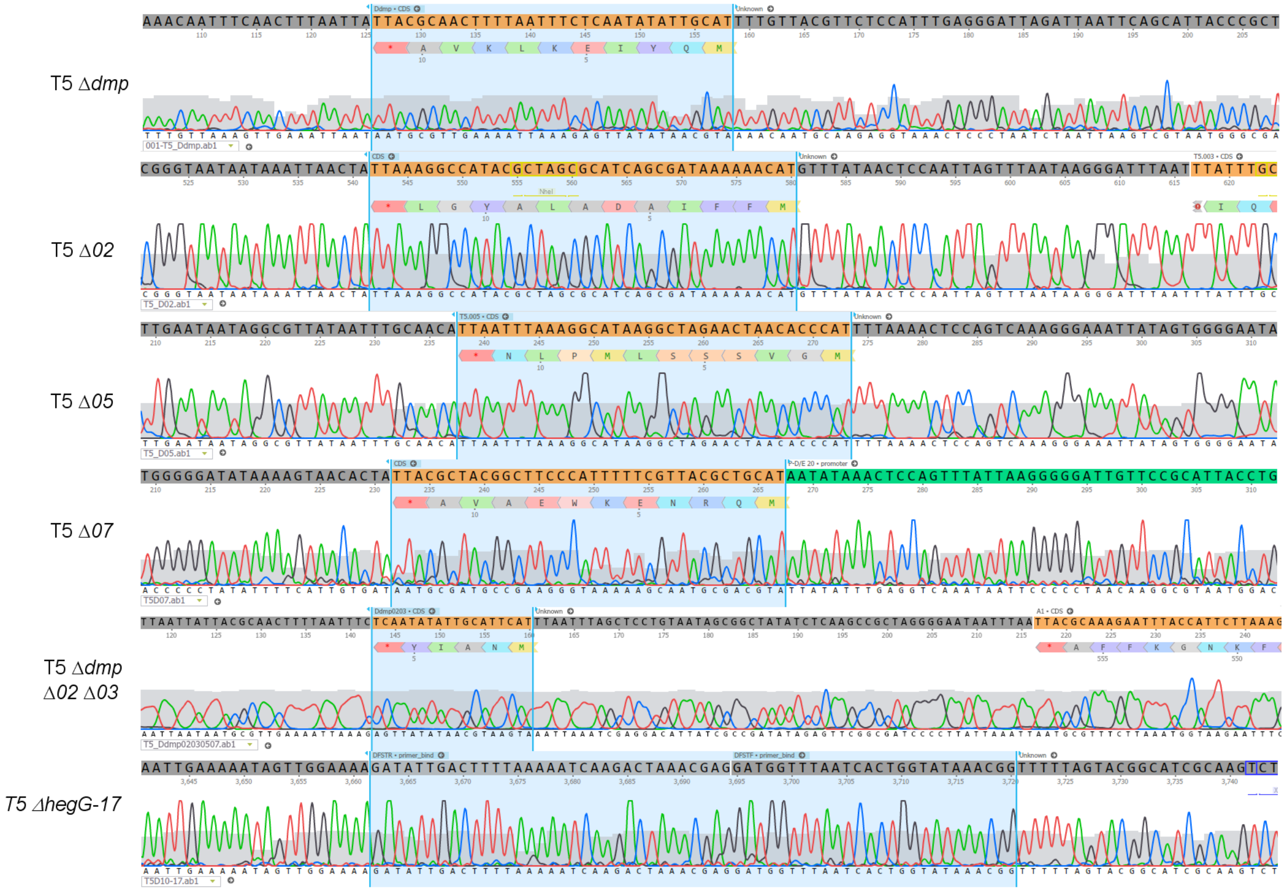


**Figure S11. Sanger sequencing from mutant strains showing the deletion sites in *dmp*, *002*, *005*, *007* and from *hegG* to *017*.**

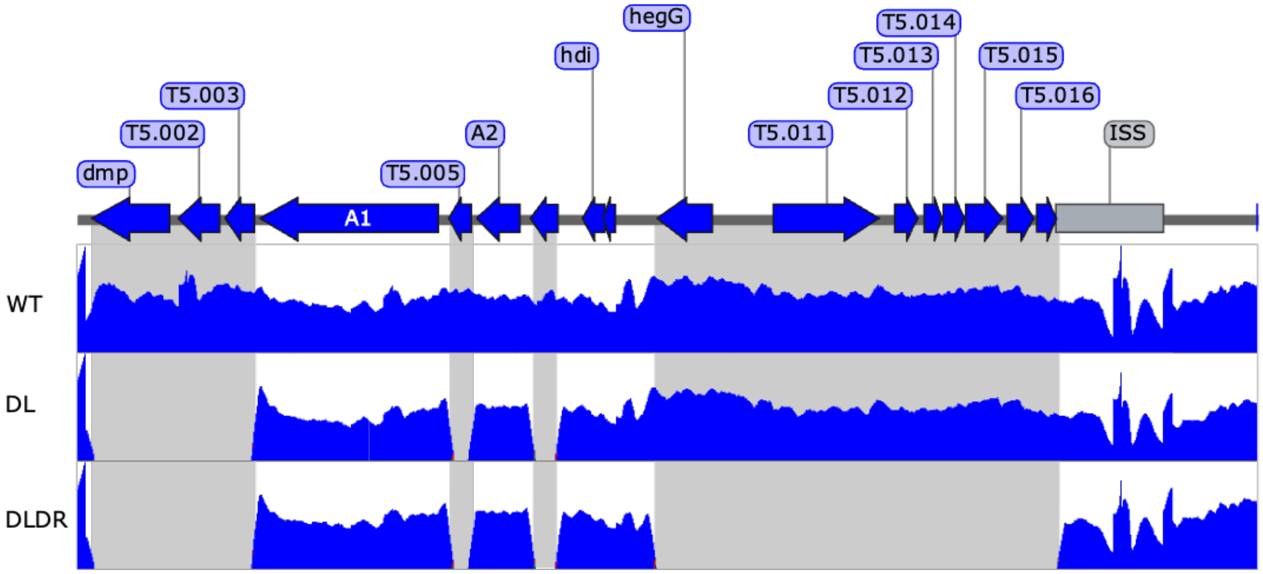


**Figure S12. Verification of mutants DL and DLDR by whole genome sequencing (WGS).**
Illumina short reads for T5 WT and mutants DL and DR were aligned to the wild-type T5 genome using Bowtie2 and Samtools with default settings. Sequencing coverage was visualized with IGV. ISS, injection stop signal. Grey shading indicates the deleted regions in phages DL and/or DLDR.

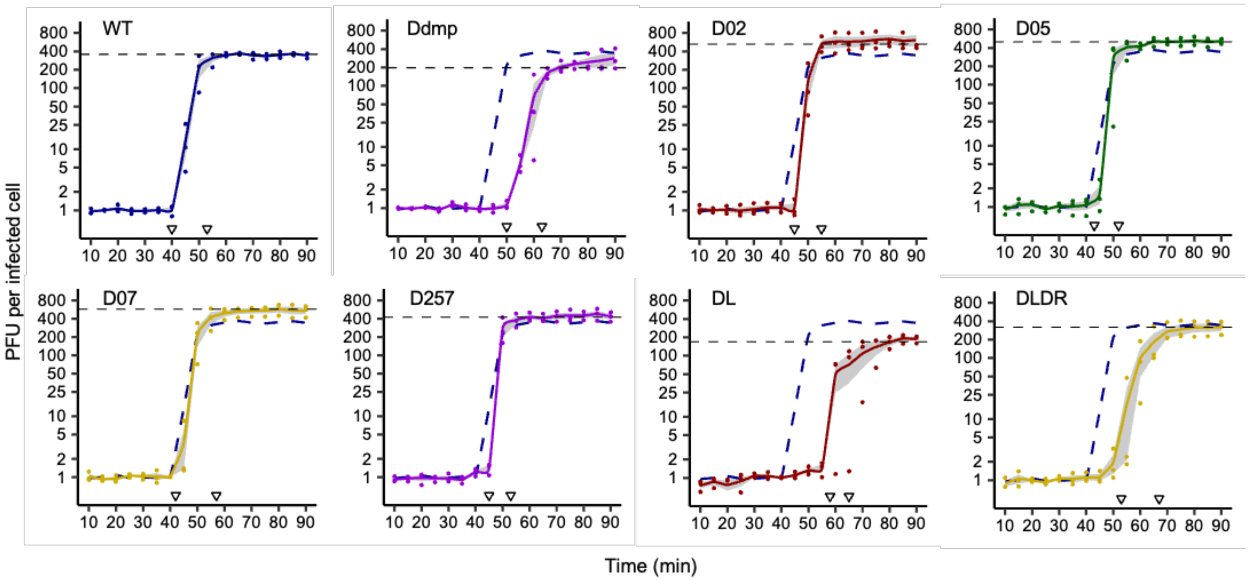


**Figure S13. One-step growth curves from mutants of T5.**
Cultures of *E. coli* strain F in exponential phase were infected at an MOI ~$10^{-3}$. Phage titers were measured every 5 minutes as described previously (46). Experiments were carried out three times. Solid lines correspond to the average of titers (dots). The surrounding grey ribbon spans one standard deviation around the average. Blue dashed curve indicates the average titers obtained for the wild-type strain. White triangles highlight the latency and the post-rise periods, whereas the horizontal dashed line indicate the burst size.

**Table S2. Pairwise comparison[a] of one-step growth parameters using ANOVA and Tukey's Honest Significant Difference post-hoc test**

| Phage | Burst size | Eclipse | Latency |
|---|---|---|---|
| Ddmp | 0.879 | **$3.87 \times 10^{-4}$** | **$6.30 \times 10^{-3}$** |
| D02 | **0.036** | 1.000 | 0.366 |
| D05 | 0.289 | 1.000 | 0.791 |
| D07 | 0.063 | 1.000 | 0.791 |
| D257 | 0.519 | 1.000 | 0.366 |
| DL | 0.700 | **$5.11 \times 10^{-9}$** | **$7.50 \times 10^{-6}$** |
| DLDR | 1.000 | **$3.87 \times 10^{-4}$** | **$3.54 \times 10^{-4}$** |

[a]Comparison between wild-type versus mutant strains regarding the Burst size, Eclipse, and Latent periods obtained from the one-step growth assays (Fig. S6). Figures in bold indicate a *p* value under 0.05.

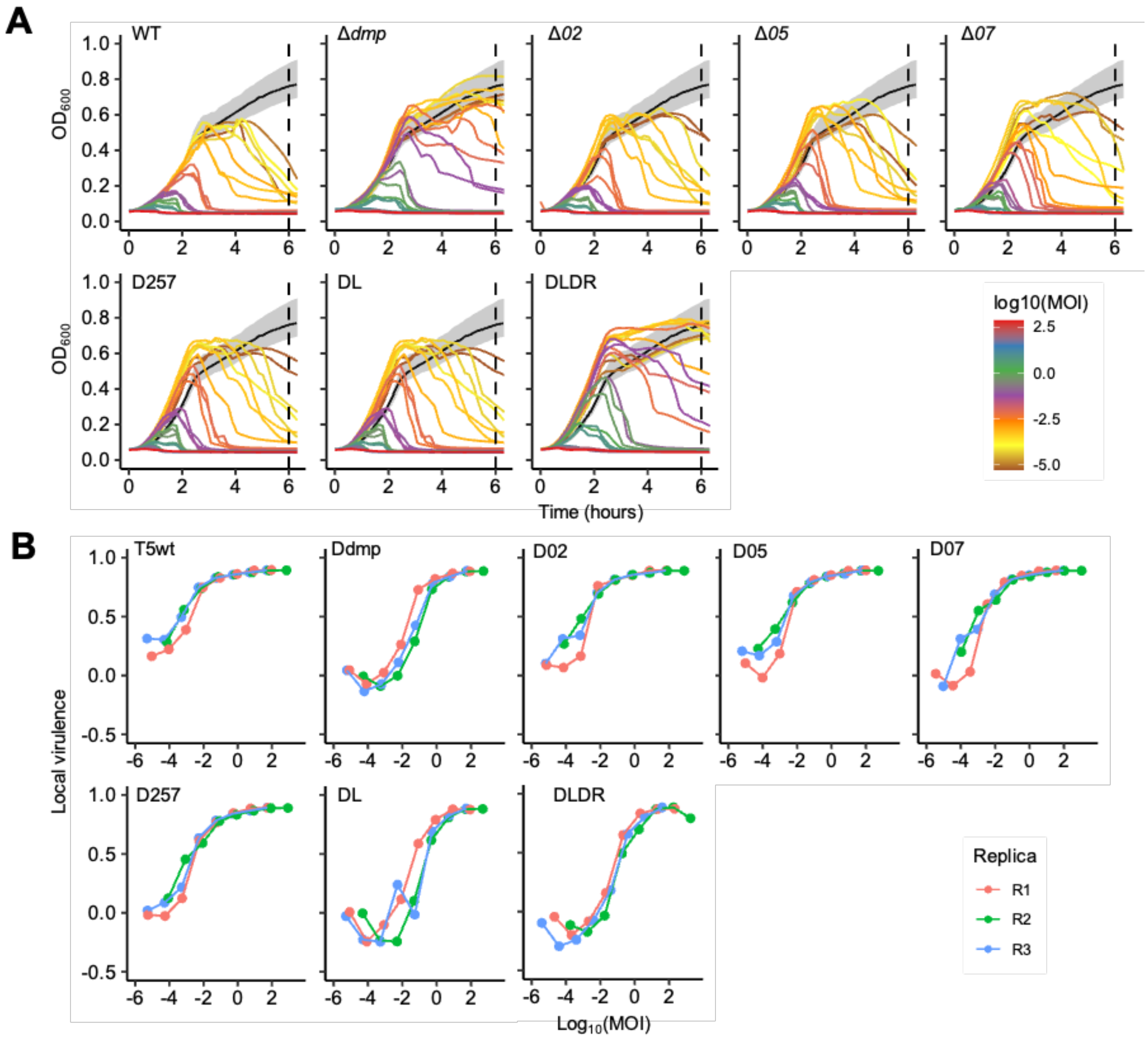


**Figure S14. Bacterial reduction and local virulence curves.**
**(A)** Bacterial reduction curves of cultures infected with T5 mutants for different multiplicities of infection (MOI). **(B)** Virulence curves represent the local virulence (calculated for each MOI using bacterial reduction curves in A) versus the logarithm of MOI. Vertical dashed lines indicate the limit of integration.

**Table S3. Dunn's post-hoc test for comparison of phage virulence [a]**

| | Phage virulence index (VI) |
|---|---|
| Kruskal-Wallis test | 0.01265 |
| T5 Ddmp | 0.0159 |
| T5 DL | **0.0040** |
| T5 DLDR | 0.0108 |

[a], comparisons listed only for $p < 0.05$ for the data presented in Fig. S7
